# Spatial regulation of Drosophila ovarian Follicle Stem Cell division rates and cell cycle transitions

**DOI:** 10.1101/2022.06.22.497017

**Authors:** David Melamed, Aaron Choi, Amy Reilein, Daniel Kalderon

## Abstract

Drosophila ovarian Follicle Stem Cells (FSCs) present a favorable paradigm for understanding how stem cell division and differentiation are balanced in communities where they can be regulated independently. Many key extracellular signals for FSCs have been identified, including inversely graded Wnt and JAK-STAT pathway activators. FSCs also exhibit interesting functional spatial heterogeneity; posterior FSCs become proliferative Follicle Cells, while anterior FSCs become quiescent Escort Cells at a much lower rate. Here, by using live imaging and FUCCI cell-cycle reporters, we measured absolute division rates and found that posterior FSCs cycle 3-4 times faster than their anterior neighbors, matching their increased differentiation rate. We also found evidence for FSC G2/M cycling restrictions and G1/S restriction that increases more anteriorly, especially beyond the FSC domain. JAK-STAT signaling promotes both transitions but graded JAK-STAT signaling alone does not explain the graded cycling of FSCs. Genetic interaction tests and FUCCI reporter assays suggest that JAK-STAT signaling acts partly through Yorkie and can largely substitute for stimulation of division by Hh signaling. PI3 kinase signaling, in contrast to Hh signaling, acts largely independently of Yorkie induction and stimulates the G2/M transition.

## Introduction

Adult stem cells must regulate their rates of division and differentiation in order to supply appropriate numbers of derivative cells without depleting the stem cell reservoir (1–3). Stem cell division and differentiation are not temporally coupled for several important highly proliferative stem cell populations, including mammalian gut and epidermal skin stem cells, as well as Drosophila FSCs, and each process can be regulated independently (4–6). In such systems of “population asymmetry” (7–9), individual stem cells and their progeny compete for survival and amplification. Consequently, genetic changes in an individual stem cell that only increase proliferation necessarily produce a competitive advantage for its descendants; such mutations may commonly be a first step towards cancer (4, 10–14). Understanding how proliferation is regulated in stem cell paradigms where differentiation is independent of division is therefore especially important for uncovering both how a stem cell pool is maintained over a lifetime and how specific genetic changes can be oncogenic.

Drosophila FSCs provide an outstanding paradigm to explore the regulation of stem cell proliferation because it has been demonstrated that altered division rates have a profound effect on stem cell survival and amplification, in agreement with theoretical expectations (4), the location and behavior of FSCs have now been clearly defined, correcting earlier misconceptions (2, 10), and important extracellular signals that regulate FSC division have been identified. Specifically, Hedgehog (Hh), JAK-STAT (Janus Kinase-Signal Transducer and Activator of Transcription) and PI3K (Phospho-Inositide 3’ Kinase) pathways were shown to stimulate FSC division and make FSCs more competitive, as were several mediators of cell cycle transitions, including CycE (CyclinE), Stg (String-ortholog of yeast Cdc25 [Cell division Control 25]) and Yki (Yorkie) (10, 15–21).

In contrast to stem cell paradigms where transitions from long-term quiescence into cycling have been the focus of investigations of proliferation (22–24), a key issue for FSCs and similar constitutively active stem cells in mammalian gut or epidermis is how the rate of division of continuously cycling stem cells is regulated. The opportunity to explore how stem cell cycling is regulated by extracellular signals is enriched in the FSC paradigm by pronounced spatial modulation; posterior FSCs cycle faster than anterior FSCs and three major signaling pathways (Hh, Wnt and JAK-STAT) are known to be graded over the FSC domain (10, 18, 25–27).

It is likely that the distribution of extracellular signals can change in response to specific stresses to allow stem cells to adapt to specific situations; indeed, the spread of Hh in the Drosophila germarium is known to depend on nutritional status (15, 16). Here we are exploring normal, optimal physiological conditions to understand how FSCs are maintained while constitutively replacing derivatives. A ring of about eight FSCs, midway along the germarium and immediately anterior to a key spatial landmark of strong Fas3 surface protein staining, directly give rise to about 5-6 proliferative FCs during each 12h cycle of egg chamber production (4, 10). Anterior and immediately adjacent to these “layer 1” FSCs are layer 2 FSCs (six, on average). These anterior FSCs and their layer 3 FSC neighbors (two, on average) directly give rise to 1-2 quiescent ECs every 12h on average. Anterior and posterior FSCs also exchange positions over time. This organization, termed “dynamic heterogeneity”, supports a stable FSC population, continuously supplying both FCs posteriorly and ECs anteriorly.

Since posterior FSCs are depleted by becoming FCs at a much higher rate (roughly four-fold) than anterior FSCs become ECs, there would be a pronounced net flow of FSCs from anterior to posterior positions if all FSCs divided at the same rate. However, posterior FSCs appear to divide faster than anterior FSCs, based on the proportion of cells in S-phase and incorporating EdU at a given time. It remains to be assessed whether this difference quantitatively matches the different rates of depletion. If the rates match, there would be little or no net AP (anterior-posterior) flux and FSCs in all locations would have equivalent life expectancies, equalizing the potential of all FSC lineages to share the collective stem cell burden. In sum, FSC proliferative control is particularly interesting because it involves regulation of both the AP pattern and absolute rates of FSC division, with a physiological outcome that dictates how stem cell function is distributed among constituents.

JAK-STAT pathway activity is stimulated by a ligand (Upd; Unpaired) released from newly-formed polar FCs and is graded across the FSC domain, with lowest activity in ECs (26). The magnitude of JAK-STAT pathway activity has a large effect on proliferation rate, indicated by EdU incorporation frequencies and FSC accumulation, and appears to be a major contributor to the AP (anterior-posterior) pattern of proliferation (18). Further understanding might benefit from better measures of cell cycling to supplement EdU studies, ascertaining which cell cycle transitions are facilitated by JAK-STAT signaling, and identifying mediators of JAK-STAT cell cycle actions. Here we employ the cell cycle reporter, FLY-FUCCI (28) and an H2B-RFP dilution strategy to those ends, and we also perform genetic epistasis tests to explore potential mediators. The effects of other pathways on FSC proliferation are less well studied and were mostly described prior to a major re-evaluation of FSC numbers, locations and behavior (10). Here we re-evaluate the responses to Hh and PI3K pathways, while exploring potential mediators. A particularly interesting potential mediator is Yorkie (Yki), a transcriptional co-activator, which can be regulated through modification of protein activity via the Hippo pathway, as well as transcriptionally (29).

## Results

### FUCCI reporters of FSC cell cycle stage

The distribution of cells among phases of the cell cycle can report which transitions appear to limit cycling in different locations and how passage through those transitions is altered by specific genetic manipulations. The FLY-FUCCI reporters link an E2F1 degron to EGFP, conferring rapid degradation in S phase, and a CycB (CyclinB) degron to mRFP1, conferring rapid degradation at the end of M phase and during G1 (28). Thus, “GFP-only” indicates cells in G1, “GFP plus RFP” indicates G2 cells (and in M phase, which can be recognized by morphology) and S phase cells ideally express only RFP. In practice, the onset of detectable RFP after the G1/S transition depends on the speed with which new protein can be made, so early S-phase cells may have no detectable GFP or RFP. Likewise, the addition of GFP to RFP at the start of G2 may have a lag, so that cells in early G2 may only have detectable RFP.

After trials with FUCCI reporters driven directly by a *ubiquitin* gene promoter or driven indirectly (*UAS-FUCCI*) by yeast GAL4 expressed from *act-GAL4*, *tj-GAL4* or *C587-GAL4* transgenes we found that *C587-GAL4* together with two copies of *UAS-FUCCI* reporters produced the strongest signals in FSCs. The *C587-GAL4* expression domain has previously been defined by driving UAS-reporter proteins that are relatively stable (GFP, RFP, β-galactosidase) and reported as being expressed strongly in ECs and declining posteriorly over the FSC domain, with some signal in the earliest FCs (10, 30). FUCCI reporters revealed a similar pattern of robust signals over the entire FSC and EC domains with a major decline in labeling at the anterior border of strong Fas3 staining, beyond which are FCs (Fig. 1). Although some Fas3-positive FCs are labeled, the demarcation of expression between FSCs and FCs is extremely useful for reporting the border between FSCs and FCs during live imaging, where Fas3 cannot be used as a precise positional marker.

**Figure 1.**
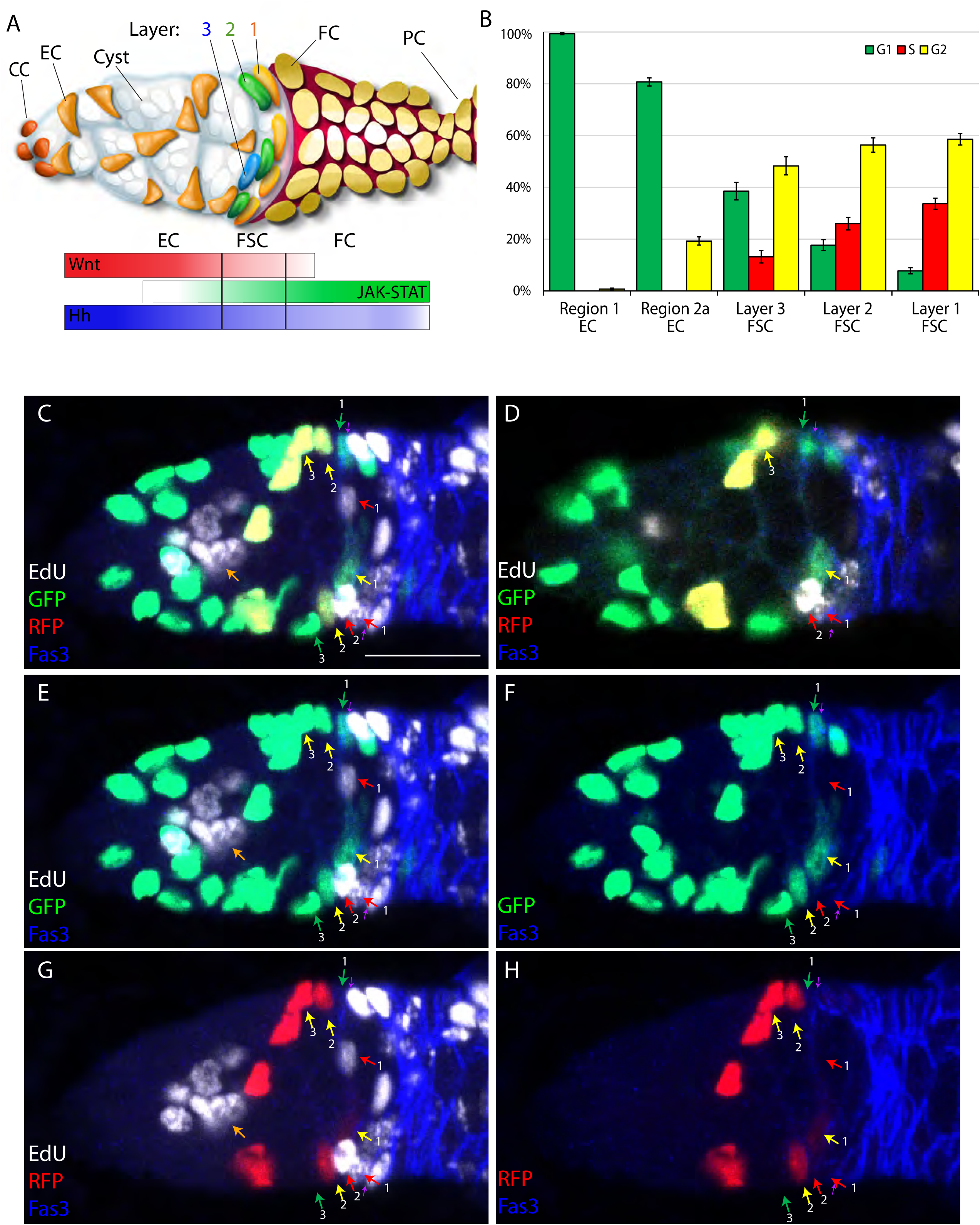
FUCCI reporter and EdU labeling to score cell cycle phase of all FSCs and ECs. (A) Cartoon representation of a germarium based on (18). Cap cells (CCs) at the anterior (left) contact germline stem cells (not highlighted), which produce cystoblast daughters that mature into 16-cell germline cysts (white) as they progress posteriorly. Quiescent Escort cells (ECs) extend processes around germline cysts and support their differentiation. Follicle Stem Cells (FSCs) occupy three AP rings (3, 2, 1) around the germarial circumference and immediately anterior to strong Fas3 expression (red) on the surface of all early Follicle Cells (FCs). FCs proliferate to form a monolayer epithelium, including specialized terminal Polar cells (PCs), which secrete the Upd ligand responsible for generating a JAK-STAT pathway gradient (green). Wnt pathway (red) and Hh pathway (blue) gradients have opposite polarity and are generated by ligands produced in CCs and ECs. (B) Percentage of cells in the designated locations in G1 (green), S (red) and G2/M (yellow), deduced from FUCCI reporters in wild-type germaria. SEM is shown from scoring 28 germaria. (C, E-H) A combination of consecutive z-sections capturing just over half of the FSCs and ECs in a single *C587>FUCCI* germarium after EdU labeling, to illustrate comprehensive scoring. Cells expressing GFP-only (green arrows), RFP-only (red arrows) or both (yellow arrows) in (C) are clarified by images showing only (E, F) GFP or only (G, H) RFP channels. Likewise, FSCs with EdU incorporation (white) are clarified by images with (C, E, F) and without (F, H) the EdU channel. All three FSCs in S-phase (red arrows) express neither GFP nor RFP. Many FCs, posterior (right) to FSCs, and a germline cyst (large clustered nuclei; orange arrow) have EdU label. The anterior border of strong Fas3 staining is indicated by purple arrows. Sample are scored by examining each z-section, as shown in (D), in order to identify the Fas3 border and consequently assign FSCs to different layers (indicated as 1, 2 or 3 for all images). Scale bar is 10μm.

Using *C587-GAL4* and FUCCI reporters together with a 1h pulse of EdU labeling immediately prior to fixation, we found that no cells with GFP-only or GFP-plus-RFP expression included EdU, consistent with assignment of all such cells to G1 and G2 (or M), respectively (Fig. 1C-H). Some EdU-positive cells included RFP but the majority were colorless, showing that the delay in accumulating RFP in S-phase is extensive. All RFP-only cells also had EdU label. Thus, S phase cells can be recognized as lacking both GFP and RFP (the majority) or expressing only RFP, and this can be confirmed in fixed tissues by EdU labeling. All other cell cycle phases are represented by a single color or color combination, and live imaging later confirmed the expected sequence of transitions. We also examined germaria stained with DAPI, instead of EdU incorporation, to visualize each somatic cell nucleus (Fig. S1). The deduced proportion of S-phase cells, counted as colorless or RFP-only, was very similar to the proportion of S-phase cells detected directly in samples labeled with EdU. Since both methods for assessing all cell cycle phases in fixed samples were robust, we combined the data from the two approaches for wild-type germaria and a few mutant genotypes, and subsequently used only DAPI-labeled samples.

### A G2/M barrier in posterior FSCs and a more prominent G1/S barrier further anterior

Consistent with earlier studies using only EdU labeling, the most posterior (layer 1) FSCs were more frequently in S-phase than their anterior neighbors (34% vs 26% for layer 2 and 13% for layer 3), while ECs were never in S-phase (Fig. 1B) (10, 18). Layer 1 FSCs were also frequently in G2 (59%), with very few in G1 (8%). Layer 2 FSCs had twice the frequency of G1 cells (18%) but the majority were still in G2 (56%), while the proportion of G1 (39%) and G2 cells (48%) were much more similar in layer 3. Region 2a ECs were mostly in G1 (81%) and the most anterior ECs (region 1) were almost entirely in G1 (99%). Clearly, ECs are restricted by an inability to transition into S-phase (and might largely be considered to be in a G0 phase), while the most posterior FSCs pass quickly through G1, with cycling apparently limited principally by a G2/M barrier. There must therefore be a robust anterior-posterior gradient, highest in posterior FSCs, of factors that stimulate the G1/S transition (or an inverse distribution of G1/S inhibitors).

We tested the effects on FUCCI reporters of altering genotypes of cells in the germarium (Figs. 2 and 3). In all cases we used GAL4-responsive transgenes, together with *C587-GAL4* and a transgene encoding temperature-sensitive GAL80. Flies were raised at the permissive temperature (18C) and shifted the restrictive temperature (29C) 3d before analysis. Trials using this protocol with reporters for signaling pathway activity, instead of FUCCI, showed significant changes by 2d and robust changes by 3d at the restrictive temperature (18). We selected the shortest effective time (3d) in order to study primary responses, prior to any systematic compensatory mechanisms, which might, for example, be triggered by accumulation or depletion of FSCs. We first explored known cell cycle mediators, before turning to the effects of different signaling pathways.

**Figure 2.**
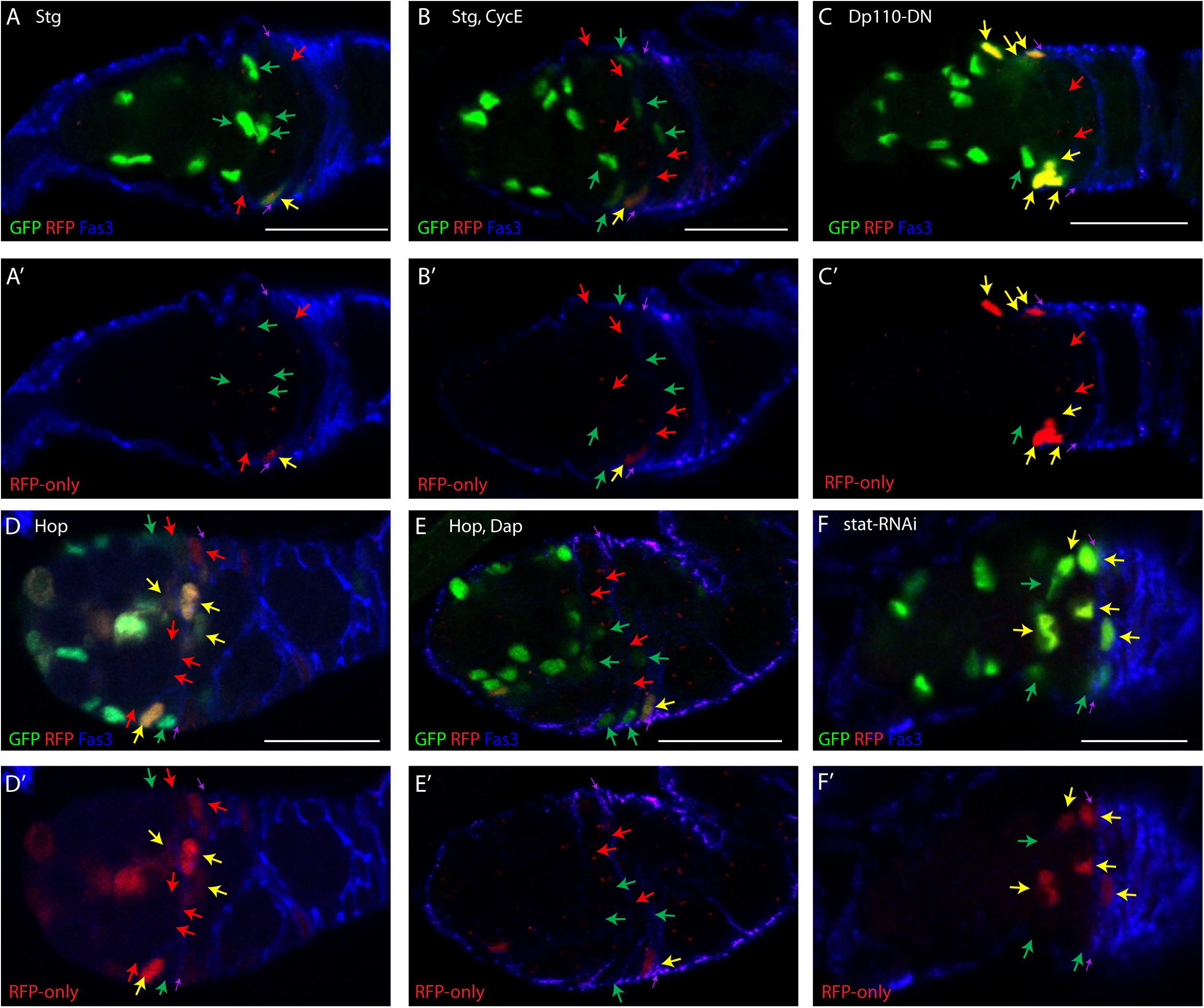
FUCCI reporter responses to key cell cycle regulators and signals. Examples of responses of FUCCI reporters to conditional expression of indicated transgenes, showing Fas3 (blue) staining to infer FSC identity and location, (A-H) GFP and RFP or (A’-H”) only RFP. All samples were also stained with DAPI to be able to count all cells, including those in S-phase (red arrows). All G1 (green arrows) and G2/M (yellow arrows) FSCs in the range of z-sections shown are indicated, as is the anterior Fas3 border (purple arrows). (A) Excess Stg increased G1 and decreased G2 FSC numbers. (B) Excess Stg together with CycE greatly increased S and decreased G2 FSC numbers. (C) Dominant-negative PI3K decreased G1 and increased G2 FSC numbers. (D) Excess JAK-STAT activity decreased G2 and increased S FSC numbers, but (E) principally increased G1 and decreased G2 FSC numbers together with excess Dacapo, Cdk2 inhibitor. (F) Decreased JAK-STAT activity increased G2 and decreased S FSC numbers. ECs were mainly in G1 in all samples but (D) and (E) illustrate conversion of some to G2 by excess JAK-STAT activity. Scale bar is 10μm.

**Figure 3.**
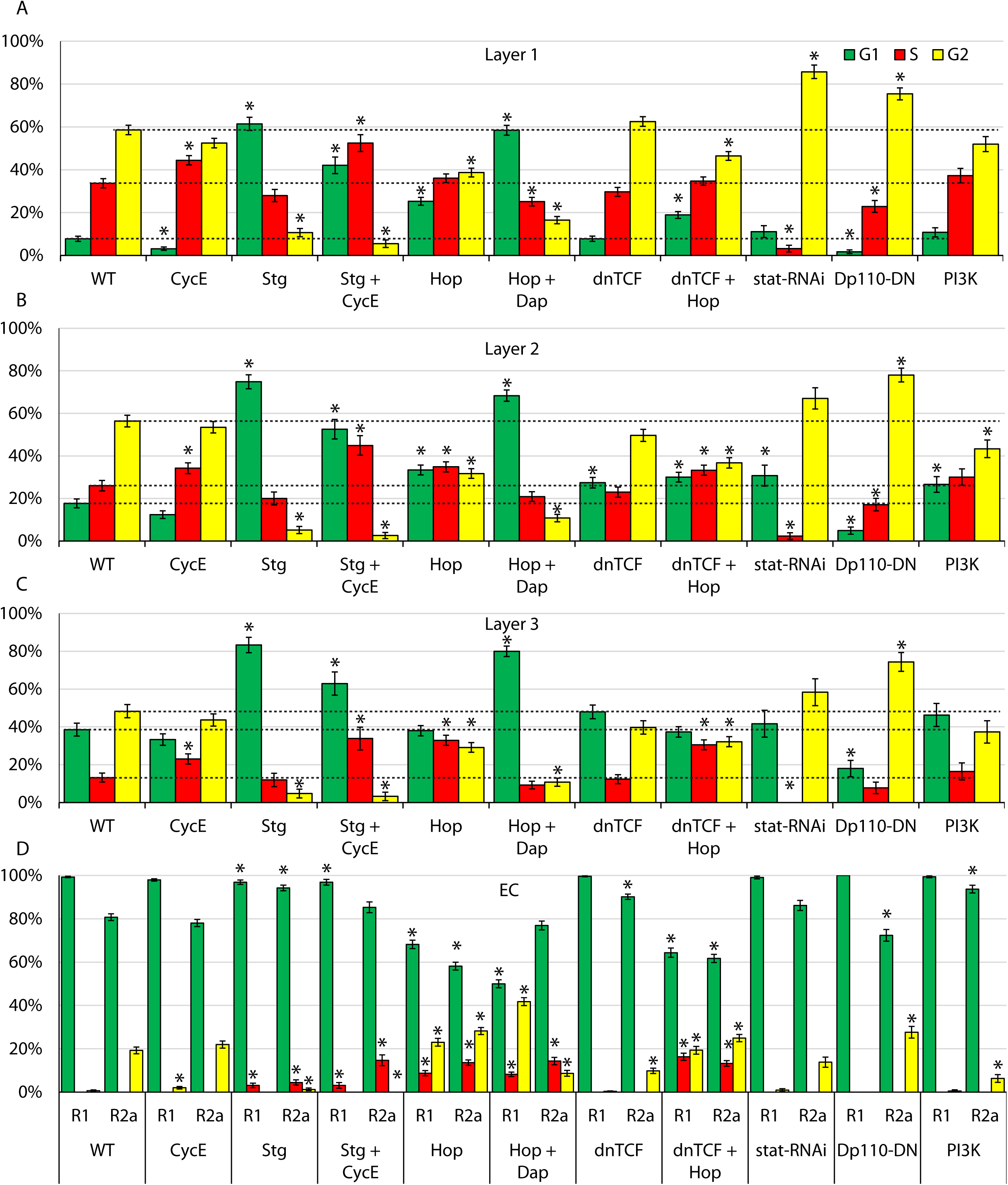
Quantitative impact of cell cycle regulators and signaling pathways on cell cycle distributions. (A-D) Percentage of (A) layer 1 FSCs, (B) layer 2 FSCs, (C) layer 3 FSCS, (D) region 2a ECs and (E) region 1 ECs in G1 (green), S (red) and G2/M (yellow), assessed by scoring FUCCI reporters for germaria expressing the indicated transgenes under *C587-GAL4* control in the presence of temperature-sensitive GAL80, after shifting to the restrictive temperature of 29C for 6d (*stat RNAi*) or 3d (all others). SEM is shown. Horizontal dotted lines indicate control values for G1, S and G2/M; significant differences for each are indicated by an asterisk (p<0.05, n-1 chi-squared test).

CycE-Cdk2 (Cyclin-dependent kinase 2) activity is a key determinant of passage into S-phase and can be regulated through the levels of CycE protein or Cdk inhibitors, such as the Dacapo (Dap) protein (31–35). In Drosophila there is a single *cycE* gene and CycE/Cdk complex, with an essential role in the cell cycle. Provision of additional CycE increased the proportion of FSCs in S phase for all three FSC layers (Fig. 3A-C), as observed previously when additional CycE, expressed from *tubulin* and *actin* gene promoters, was limited to clones of cells derived from a single FSC (18). The proportion of cells in G1 was diminished in layer 1 FSCs (from 7.8% to 3.1%), as might be expected for promoting the G1/S transition (Fig. 4). The proportional reduction in G1 cells in layer 2 (17.7% to 12.4%) and layer 3 (38.5% to 33.3%) were lower, with a notably high residue of G1 layer 3 FSCs. There were also small reductions of the proportions of cells in G2 for all FSC layers. The proportion of G1 cells in region 2a (78%) and region 1 ECs (98%) remained substantially unchanged. Thus, although excess CycE appears to promote G1/S transitions in the most posterior FSCs, it is surprisingly ineffective in more anterior FSCs and especially in ECs. This suggests that the G1/S transition in ECs and anterior FSCs is restricted by a factor other than CycE-Cdk2 activity or that the deficit in CycE/Cdk2 activity is too large to be reversed by the strategy we used.

**Figure 4.**
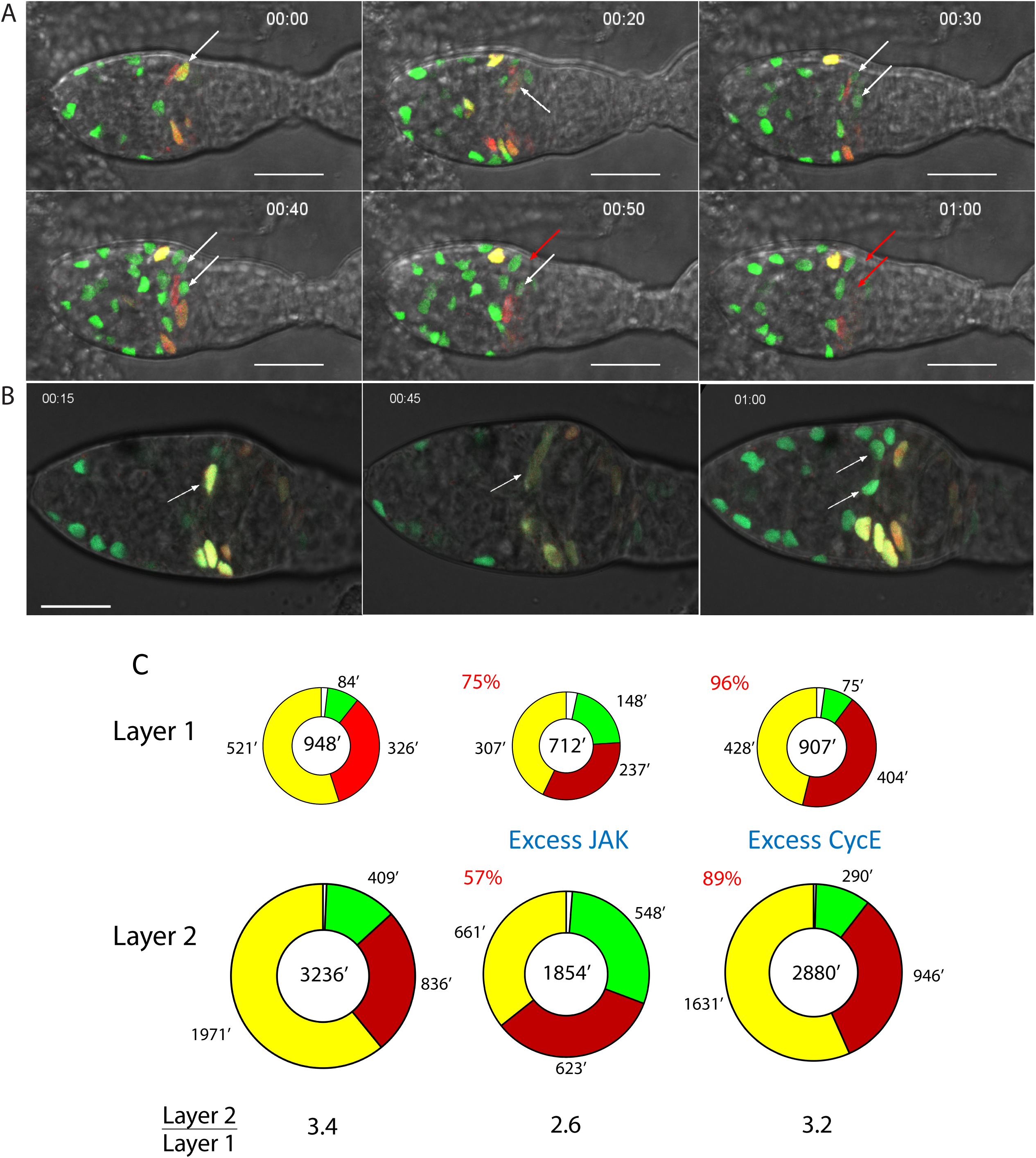
Live FUCCI reporter imaging shows cell cycle transitions and phase durations. (A, B) Time-stamped frames from live imaging of (A) control and (B) *C587>Hop* germaria. (A) A yellow G2 cell (white arrow) changes to M-phase morphology at 20min, producing two G1 daughters (white arrows) with low GFP (no RFP) signal at 30min, strengthening by 40min. One daughter (red arrow) has lost GFP, indicating S-phase (red arrow) at 50min. The second daughter lost GFP, entering S-phase at 60min. The highlighted cell and its daughters are scored as layer 1 FSCs because they are at the posterior margin of strong *C587-GAL4* expression driving the FUCCI reporter transgenes. (B) A yellow G2 cell (white arrow) changes to M-phase morphology at 45min, and then produced two G1 (green) daughters (white arrows) by 1h. The highlighted cells are two diameters away from the posterior edge of *C587-GAL4* expression, indicating that they are layer 3 FSCs. Cell cycle transitions for layer 3 FSCs were observed only for cells with increased JAK-STAT activity, consistent with slow cycling of anterior FSCs resulting partly from insufficient JAK-STAT activity. Scale bar is 10μm. (C) Summary of calculated duration of G1 (green), S (red), G2 (yellow) and M (white) phases of the cell cycle from the sum of all live imaging assays for control (left), *C587>Hop* (middle) and *C587>CycE* (right) germaria. For each genotype, the ratio of the whole cell cycle length for layer 2 relative to layer 1 FSCs is shown. The lengths of cell cycles as a percentage of controls are shown in red for excess JAK and CycE.

The activity of Stg is critical for G2/M passage and is often regulated transcriptionally (36–40). Additional Stg expression dramatically reduced the proportion of FSCs in G2 in all layers (59% to 11%, 56% to 5%, 48% to 5%) (Fig. 2A and Fig. 3), consistent with earlier studies that showed a large increase in the proportion of cells positive for the M-phase marker, phosphor-histone H3 (19). There was no increase in the proportion of FSCs in S phase, with the cells accelerated through G2 apparently accumulating in G1 (61% in layer 1, 75% in layer 2, 83% in layer 3). Stg overexpression was found previously to shorten G2 and extend G1 in Drosophila wing disc cells; further study suggested that the extension of G1 involved the action of E2F1, which mediated compensatory feedback mechanisms between G1 and G2 when either was artificially altered (41). When both additional Stg and CycE were expressed in ECs and FSCs, the fraction of FSCs in G2 was also very low (<6%) in all layers but the proportion of cells in S phase was substantially higher in all layers than in controls (52% vs 34%, 45% vs 26%, 34% vs 13%) (Fig. 2B and Fig. 3) or in cells expressing only excess Stg. Thus, in Drosophila FSCs, excess Stg potently promoted mitosis and caused the majority of FSCs to accumulate in G1. Adding excess CycE significantly reduced this G1 accumulation but the proportion of FSCs in G1 was still much higher than normal for all FSC layers (42% vs 8%, 53% vs 18%, 63% vs 39%). The combination of excess Stg and excess CycE appears to accelerate FSC cycling substantially in all layers judged by the S-phase index (which has the important caveat that the length of S phase is unknown). A distinct gradient of increasing S-phase proportions from anterior to posterior remained, again suggesting a spatially graded G1/S restriction that is not readily overcome by excess CycE.

Excess Stg virtually eliminated the (19%) fraction of region 2a ECs normally in G2 (Fig. 3D). Surprisingly, a small fraction of region 2a (4%) and region 1 ECs (3%) were found in S-phase rather than G1. These fractions were increased to 15% and 12%, respectively, when excess CycE was co-expressed. Thus, besides efficiently stimulating mitosis in ECs, excess Stg appears to result in facilitated G1/S passage, especially in combination with excess CycE.

### JAK-STAT signaling promotes G1/S and G2/M transitions

Excess Hop (Drosophila JAK) substantially decreased the fraction of FSCs in G2 for all layers (59% to 39%, 56% to 32%, 48% to 29%) (Fig. 2D and Fig. 3A-C). In contrast to the effects of additional Stg, the fraction of FSCs in S phase was significantly increased, especially in layer 2 (26% to 35%) and layer 3 (13% to 33%), where *C587-GAL4* is expressed most strongly. The comparison to the effects of Stg indicates that excess JAK-STAT signaling strongly promotes the G1/S transition as well as mitosis, even though the proportion of FSCs in G1 is significantly higher than in normal cells in layer 1 (25% vs 8%) and layer 2 (33% vs 18%). In region 2a and region 1 ECs, excess JAK-STAT activity significantly reduced the fraction of cells in G1 (81% to 58%, 99% to 68%), increased the fraction in S-phase (from 0 to 14% and 9%, respectively) and in G2 (19% to 28%, 1% to 23%). Thus, in ECs the most prominent action of increased JAK-STAT is to stimulate G1/S passage, leading to significant accumulation of cells in both S and G2 phases.

By comparison to the effects of additional Stg or Stg plus CycE, it appears that increased JAK-STAT stimulates the G1/S transition more potently than G2/M passage in ECs, whereas the converse appears to be true for FSCs. A JAK-STAT pathway reporter was used previously to demonstrate that pathway activity is nearly uniform across all FSCs under the conditions of this experiment (at about 140% of normal layer 1 FSC levels) and slightly lower in ECs (only marginally higher than in normal layer 1 FSCs). The different potency of excess JAK-STAT in different locations therefore presumably reflects constraints normally imposed by spatially restricted factors other than JAK-STAT ligand.

When the CycE/Cdk2 inhibitor Dacapo (Dap) was co-expressed with *UAS-Hop* the proportion of cells in G2 was decreased more than for *UAS-Hop* alone in each FSC layer (59% to 16%, 56% to 11%, 48% to 11%) but the fraction in S-phase was in each case decreased (34% to 25%, 26% to 21%, 13% to 9%), with the majority of FSCs accumulating in G1 (58%, 68%, 80%) (Fig. 2E and Fig. 3A-C). Thus, Dap appears to override the G1/S stimulation provided by excess JAK-STAT pathway activity in FSCs. Dap did not, however, reduce the fraction of ECs driven into S-phase by *UAS-Hop*, even though the distribution between G1 and G2 was somewhat altered. Thus, promotion of G1/S transitions in FSCs by increased JAK-STAT pathway activity was phenocopied by excess CycE and inhibited by a CycE/Cdk2 inhibitor, whereas both excess CycE and Dap were without effect in ECs, suggesting that excess JAK-STAT may use different mechanisms to promote G1/S transitions in FSCs and ECs (the pathway normally has only low activity in ECs). The strong effect of increased JAK-STAT pathway activity on exit from G2 in all FSC layers was even more obvious in the presence of excess Dap.

Previous investigation of Wnt pathway reduction or elimination in FSC lineages showed important roles in guiding FSC locations and differentiation, but no significant changes in either the magnitude or pattern of cell division, inferred from EdU incorporation (18). Increased Wnt pathway activity in FSC lineages, to levels estimated at twice the physiological maximum in the germarium, severely reduced EdU incorporation but this was overridden by excess JAK-STAT pathway activity (18). Also, Wnt pathway reduction promoted EdU incorporation, in concert with excess CycE, when JAK-STAT activity was eliminated (18). Thus, Wnt pathway activity appears to have the potential to reduce FSC cycling but that potential was blocked by JAK-STAT pathway activity when assayed simply by EdU incorporation in FSC lineages.

Here, we used expression of dominant-negative TCF (dnTCF; “T-Cell Factor”, sole transcriptional effector of Drosophila Wnt signaling) to reduce Wnt signaling over the entire EC/FSC domain. Residual levels of signaling over EC and FSC regions was previously shown, using a *Fz3-RFP* reporter, to be lower than normally found in layer 1 FSCs, and to be spatially uniform, in contrast to the normal strong decline from anterior to posterior (18). Only small changes in FUCCI reporters were observed in FSCs, with slightly more layer 2 FSCs in G1 as the only notable change (Fig. 3A-C). The distribution of FSC cell cycle phases in the presence of excess JAK-STAT pathway activity was virtually unchanged when Wnt signaling was also inhibited (*dnTCF*+*UAS-Hop*; Fig. 3A-C). These results confirm prior conclusions that Wnt signaling under normal conditions, and in the presence of excess JAK-STAT activity, has very little effect on FSC cycling. In ECs, Wnt inhibition modestly reduced the frequency of region 2a G2 cells (from 19% to 10%) and all remaining cells were in G1 (Fig. 3D). In the presence of excess JAK-STAT, Wnt inhibition provoked no significant changes in the cell cycle profile. Thus, even though Wnt signaling is highest in ECs and JAK-STAT signaling is low or absent in most ECs, the prominent G1/S barrier in these cells does not depend on Wnt pathway activity.

Depletion of gene products using GAL4-responsive *UAS-RNAi* transgenes can take longer than increasing protein activities by overexpression. Expression of *stat RNAi* had only small effects on FUCCI reporters after 3d (data not shown) but robust effects by 6d, so we conducted FUCCI analysis at 6d. In layer 1 FSCs, the proportion of cells in S-phase was dramatically reduced (34% to 3%), with many additional cells accumulating in G2 (59% to 86%), indicating a major deficit in entering mitosis (Fig. 2F and Fig. 3A-C). The fraction of cells in S-phase was also greatly reduced in layer 2 (26% to 2%) and layer 3 (13% to 0). In these more anterior locations the increased number of cells in other phases were more equally distributed between G1 (18% to 31%, 39% to 42%) and G2 (56% to 67%, 48% to 58%), suggesting that the loss of JAK-STAT reduced the frequency of both G1/S and G2/M transitions. These observations are consistent with deductions from responses to increased JAK-STAT activity, indicating JAK-STAT promotion of G2/M transitions, most prominently in posterior FSCs, and also G1/S transitions, more prominent in anterior FSCs. The FUCCI profile of ECs, where JAK-STAT activity is normally very low, was barely altered by loss of STAT (Fig. 3D).

### PI3K pathway promotes entry into mitosis throughout the FSC domain

In Drosophila, insulin-like growth factor binding to the insulin receptor is transmitted via the insulin-like receptor substrates, Chico and Lnk, to activate PI3K (42–45). As in mammals, downstream responses to PI3K include Tor complex-mediated translational and metabolic changes and FoxO-dependent transcriptional changes, generally integrated to promote cell growth and proliferation. Unlike in mammals, activation of other tyrosine kinase receptors principally activates Ras/MAPK signaling, without concerted PI3K pathway activation. When PI3K pathway activity was inhibited by expression of a dominant-negative version of the catalytic subunit, Dp110 (aka PI3K92E, here “PI3K”) the fraction of FSCs in G2 was significantly increased in all layers (Fig. 2C and Fig. 3A-C). The increase in layer 1 (from 59% to 75%) was less than due to STAT loss (86%) but the increase in layers 2 and 3 were greater than observed for STAT loss (56% to 78% vs 67%, 48% to 74% vs 58%). Thus, PI3K pathway activity appears normally to promote exit from G2 in FSCs and shares this responsibility with JAK-STAT signaling, with a greater role in more anterior cells where JAK-STAT pathway activity is normally lower. The AP profile of PI3K pathway activity has not been determined but these results suggest it is unlikely that the pathway is higher in more posterior cells, like the JAK-STAT pathway. The proportion of FSCs in G1 was reduced by more than the proportion in S-phase in all layers when PI3K pathway activity was reduced (8% to 2% vs 34% to 23%, 18% to 5% vs 26% to 17%, 39% to 18% vs 13% to 8%), suggesting that the pathway does not normally play a prominent role in G1 exit. By contrast, loss of JAK-STAT was associated with a much greater reduction in S-phase cells, implying an important role in promoting the G1/S transition.

The effects of increased expression of the PI3K catalytic subunit (using “*UAS-PI3K*”) were smaller (only changes in layer 2 FSCs were statistically significant) and complementary, reducing the fraction of FSCs in G2 in all layers (59% to 52%, 56% to 43%, 48% to 37%), and increasing the fraction of cells in G1 (8% to 11%, 18% to 27%, 39% to 46%) and S-phase (34% to 37%, 26% to 30%, 13% to 16%) to similar degrees (Fig. 3). These results suggest some potential for excess PI3K to promote G2 exit and perhaps G1 exit (because S-phase frequency was marginally increased, whereas excess Stg, which also primarily stimulates G2 exit, reduced the fraction of cells in S-phase). These effects were much smaller than observed for excess JAK-STAT, especially for anterior FSCs, where only excess JAK-STAT promoted an increase in S-phase cells. Thus, the PI3K pathway appears to promote the G2/M transition throughout the FSC domain and may have some, lesser potential to facilitate G1 exit. These properties appear to extend to the EC domain, where increased PI3K activity reduced region 2a cell G2 proportions (from 19% to 6%) but still with no cells in S-phase. PI3K pathway inhibition increased the fraction of region 2a G2 ECs to 29%; all region 1 ECs remained in G1 in both genetic conditions.

### Live FUCCI Imaging to infer cell cycle times

The proportion of cells of a certain type, or in a restricted location, that label with the mitotic marker, phospho-histone H3, or incorporate EdU as an indicator of S-phase, is often used as a proxy to estimate either the proportion of cycling cells or the average rate of cell cycling. The fraction of cells in M phase or S phase under different conditions will, however, only provide a measure of the relative cell cycle times if the length of that phase remains constant. While regulation at G1/S and G2/M transitions is common, there is no guarantee that the length of M or S phase is constant for a given cell type under different conditions or in different locations. We therefore used FUCCI reporters, together with a previously developed system for live imaging of germaria freshly dissected from flies and embedded in Matrigel (10, 46), to measure absolute cell cycle times.

We deliberately imaged for only short periods of time (generally close to 2h; average 148min) in order that the behavior of cells more likely reflects normal *in vivo* behavior, even though viable, active germaria can be imaged for as long as 8-12h (10, 46). The pattern of *C587-GAL4* expression, mostly terminating with layer 1 FSCs, allowed us to identify layer 1, layer 2 and layer 3 FSCs even though there was no Fas3 staining landmark and germaria consistently moved during imaging. Imaging each z-section every 20 mins or less allowed us to track each labeled cell with confidence. Some cells moved out of the z-section range during imaging, while other cells in the same germarium were imaged for longer. We observed each type of expected transition: loss of GFP, indicating a G1/S transition, entry of G2 cells into mitosis, which was generally observed morphologically in only a single frame, followed by loss of RFP to leave two GFP-only nuclei of daughters in G1 (Fig. 4A, B). Occasionally, an S to G2 phase transition was observed with GFP signal added to RFP, but the fate of cells initially in S-phase was not systematically tracked because the majority of S-phase cells have no detectable RFP or GFP. Thus, quantitation focused on cells in G1 (GFP only), G2 (GFP plus RFP) and mitosis. Generally, 3-8 FSCs were observed to transition from one phase to another in a single germarium. We therefore excluded data where no transitions were observed; often, such germaria exhibited minimal cell movements in contrast to apparently healthy germaria.

We reasoned that each cell in a specific phase (say, G2) was captured at an arbitrary time within that phase at the start of imaging, so if we add together the total time spent by all cells of a specific type (say, layer 1 FSCs) in that phase and divided by the number of transitions observed out of that phase we could infer the average phase duration. Collectively, we followed 90 layer 1 FSCs among fifteen germaria, spending a total of 8343 mins in G2 with 16 transitions out of G2 (to give an estimate of G2 as 521 mins), and 1681 mins in G1 with 20 transitions to S phase (to give an estimate of G1 as 84 mins). Similarly, the average time spent in M-phase was 17 mins (11 transitions through mitosis). From fixed data, time spent in S phase as a fraction of G1+G2 was 35/65, so the estimated duration of S phase was (521+84) x 35/65 = 326 mins and the total cell cycle time was 948 mins. The duration of G1 (9%) and G2/M (56%) as a fraction of the whole cell cycle of layer 1 FSCs from live imaging matched fixed image FUCCI data quite well (6% G1, 59% G2/M), providing some assurance of correct cell location identification and the absence of a major change induced by the live imaging procedure. The total number of cells followed in different layers (90 in layer 1= 50%, 71 in layer 2=39%, 20 in layer 3 =11%) also roughly matched the proportion of FSCs known to be in layers 1 (50%), 2 (37%) and 3 (13%), further confirming appropriate cell location identification.

An analogous strategy was used to estimate average layer 2 FSC cell cycle phase times from 71 tracked cells for G1 (409 mins), S (836 mins), G2 (1971 mins), and M (20 mins), summing to 3236 mins (Fig. 4C). Only four transitions from G2, and three from G1 were observed in total, so cell cycle time estimates have a much larger margin of error than for layer 1 FSCs. Nevertheless, it is clear that layer 1 FSCs cycle much faster than layer 2 FSCs, estimated as a factor of more than 3-fold (3236/948= 3.41) even though EdU incorporation indices had a much lower ratio (0.35/0.26 = 1.35). The substantial difference is because the length of S-phase, deduced from live imaging, appears to be much longer in layer 2 (836 min) than in layer 1 FSCs (326 min). This inference was also supported by direct observation. When cells transitioned from G1 they were observed to lose GFP but not to gain RFP within the subsequent observation period (ranging from 15 to 200 min; average 114 min, 22 examples), consistent with the expectation that there is a delay before sufficient RFP accumulates to visible levels. Thus, early S phase cells are colorless and late S phase cells have RFP. EdU labeling shows that layer 1 FSCs are more frequently in S-phase than layer 2 FSCs. Despite this, among 90 layer 1 FSCs, an RFP-only cell was seen for a total of only 80 mins, whereas RFP-only layer 2 FSCs (from a smaller sample of 71) were seen for a total of 789 mins, consistent with a considerably longer S-phase. None of the 20 layer 3 FSCs tracked underwent a FUCCI reporter change, consistent with the expected slow cycling of those cells but providing no quantitative data with regard to cell cycle times.

The deduced cell cycle times for FSCs in different locations can be usefully compared to indirect deductions from cell lineage studies. In an extensive study of the cells resulting from marked FSCs over a 3d period, it was ascertained that about four FCs are produced for every EC produced (4). It has also been shown that 5-6 FCs are produced per egg chamber budding cycle of about 12h, that FCs derive directly from layer 1 FSCs and ECs derive directly from anterior FSCs (4, 10). Although FSCs can exchange AP locations, if there were no net flow in the AP direction, production of 5-6 (say 5.5) FCs from 8 layer 1 FSCs (without depletion) in 12h would require an average cell cycle time of 1047 mins (12 x 8/5.5 = 17.45h). Live imaging studies revealed a cell cycle time of 931 mins. For EC production at ¼ the rate of FCs to be sustained by division of only six layer 2 FSCs (the few FSCs in layer 3 divide much more slowly), those FSCs should have a cell cycle time three-fold (4 x 6/8) greater than for layer 1 FSCs. Live imaging studies deduced a ratio of 3.4. Thus, the cell cycle durations measured by live imaging support a scenario where FC production can be supported by layer 1 FSC divisions and EC production can be supported by anterior FSC divisions without any net flow of FSCs from anterior to posterior or vice versa.

### Cell cycle times for increased CycE and increased JAK-STAT signaling

Live imaging and tracking were also performed for germaria where CycE or Hop was overexpressed, using the same genetic conditions as for fixed-sample FUCCI studies, 3d after shifting to the restrictive temperature. For excess CycE, tracking 36 layer 1 FSCs yielded deductions of a G1 phase of 75 mins, S phase of 408 mins, G2 of 404 mins, and M phase of 20 min, summing to a total cell cycle time that is 96% (907/948) of controls (Fig. 4C). The EdU index was 1.3-fold (46%/35%) higher than for wild-type layer 1 FSCs but most of this increase can be attributed to a longer S-phase (408/326= 1.25). Excess CycE was found to shorten G1 but extend S-phase in wing disc cells (41). Thus, live FUCCI imaging suggests that excess CycE accelerates layer 1 FSC cycling, but much more modestly than estimated from examining EdU incorporation indices. From 33 layer 2 FSCs, G1 duration was deduced to be 290 mins, S was 946 mins, G2 was 1631 mins, and M was 13 mins, for a total of 2880 mins, which is (2880/3236) 89% of the control cell cycle time. The EdU index increase from fixed imaging was by a factor of 1.3 (33%/26%), while S-phase was deduced to increase by a factor of 1.13 (946/ 836)). Thus, excess CycE appears to speed cycling in layer 2 FSCs by about 10% and also to lengthen S-phase by a similar factor, but all layer 2 measurements from live imaging have substantial error margins.

Excess JAK-STAT resulted in deduced phase durations of G1: 148 mins, S: 237 mins, G2: 303 mins, and M: 24 mins among 65 layer 1 FSCs for a total cell cycle time of 712 mins, which is 75% (712/948) of controls (Fig. 4B, C). For 39 layer 2 FSCs, deduced phase durations were G1: 548 mins, S: 623 mins, G2: 661 mins, and M: 22 mins, for a total cell cycle time of 1854 mins, which is 57% (1854/3236) of controls. The reduction of cell cycle time in layer 1 FSCs is much greater than for CycE overexpression even though EdU indexes were similar because S-phase duration was increased only by CycE (it was decreased by JAK-STAT). The difference was even greater for layer 2 FSCs (where the EdU index is also higher for excess JAK-STAT vs CycE). These results are consistent with the significantly greater amplification of FSCs induced by increased JAK-STAT activity, despite an increased frequency of FSC to FC conversion (18). The number of new cells produced every 12h when JAK-STAT pathway activity is increased is expected to be 8.1 (8 x 720/712) for 8 layer 1 FSCs (compared to 6.1 (8 x 720/948) for wild-type) and 2.3 (6 x 720/1854) for 6 layer 2 FSCs (compared to 1.3 (6 x 720/3236) for wild-type), and there may also be FSCs produced from layer 3 FSCs. Total FSC production per 12h budding cycle is therefore estimated to be at least 10.4 cells compared to 7.4 for controls, an increase of more than 40%.

The fractional decrease in cell cycle time induced by excess JAK was greater for layer 2 than layer 1 FSCs, consistent with the anterior bias of *C587-GAL4* expression and the higher levels of endogenous JAK-STAT activity in layer 1 FSCs. However, the deduced cell cycle time for layer 1 FSCs (712 mins) was still less than half that of layer 2 FSCs (1854 mins), even though JAK-STAT activity was roughly spatially uniform. We previously noted that the EdU indices in the two layers were identical (33%) under those circumstances (18). The discrepancy between the two measurements is because the length of S-phase remains substantially different between layers 1 and 2, even when JAK-STAT pathway activity has been equalized. Thus, less frequent entry and longer passage in layer 2 FSCs happen to balance exactly the more frequent and shorter passage through S-phase in layer 1 FSCs to give the same EdU index.

In summary, live imaging of FUCCI reporters has allowed deduction of the absolute timing of different cell cycle phases (Fig. 4C). The results revealed that the length of S-phase was not constant among all FSC locations and genotypes investigated, with the consequence that EdU index cannot reliably be used to infer cell division rates quantitatively. This limitation is most severe when comparing FSCs in different layers because S-phase is much longer for layer 2 (836 min) and layer 1 FSCs (326 min). The shortcomings of EdU index comparisons between layers were clearly exposed for conditions that produced roughly equal JAK-STAT signaling over the whole FSC domain; layer 1 FSC cell cycle time (712 min) was still much shorter than for layer 2 (1854 min), even though both layers had the same EdU index. Comparisons of different genotypes for a single FSC layer also showed that the length of S-phase was not constant, but here the variations observed were much smaller than between layers. Thus, S-phase in layer 1 FSCs was lengthened (from 326 min to 408 min) by increased CycE, and shortened (to 237 min) by increased JAK-STAT activity, with the consequence that EdU measures under-estimated the significantly greater potency of JAK-STAT in stimulating FSC division. Nevertheless, both genetic changes were found to increase division of layer 1 and layer 2 FSCs using EdU measurements and FUCCI live imaging, suggesting that EdU measurements are likely to provide a useful qualitative indication of division rate changes due to various other genotypes within a specific FSC layer.

### FSC division rate tracked by H2B-RFP dilution

We additionally aimed to measure FSC division frequency by using dilution of stable H2B-RFP protein. We mobilized a *UAS-H2B-RFP* P-element insertion and identified a third chromosome insertion with particularly strong expression when driven by *act-GAL4*. In animals also containing a temperature-sensitive *GAL80* transgene, we found that there was little H2B-RFP expression unless animals were moved from 18C to 29C, and we explored various protocols for initiating H2B-RFP expression. Similar results were obtained for any incubation at 29C longer than about 48h, whether during adulthood, pupal stages or throughout post-embryonic development. Although it might be expected that H2B-RFP is synthesized and incorporated into chromatin more efficiently in dividing cells, we observed efficient labeling of quiescent adult ECs even over short time periods. Overall, the pattern of labeling intensities among different cells were very similar to those of GFP from a *UAS-GFP* transgene also present. Under all tested conditions we observed uneven H2B-RFP (and GFP) intensities among germarial cells. FC labeling was always weaker than for FSCs and ECs, while r2a EC signals were particularly strong (Fig. 5A). We then undertook chase experiments after 4d at 29C, looking for H2B-RFP dilution. We aimed for semi-quantitative analyses, rather than being able to deduce the literal number of divisions over a chase period, because there was also some variation among H2B-RFP intensity for cells in a given AP location prior to the chase period.

**Figure 5.**
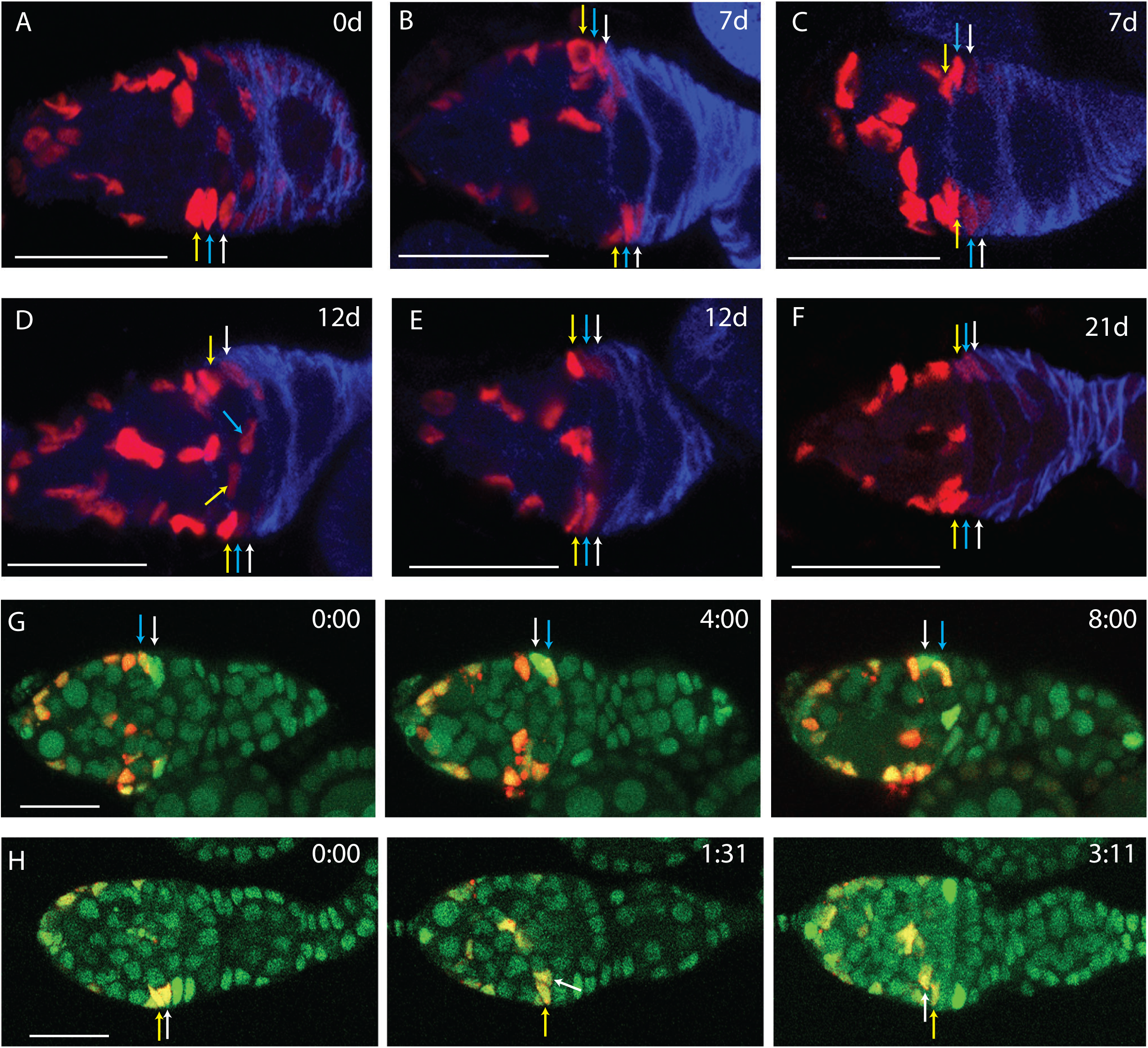
H2B-RFP dilution shows AP division gradient and exchange of FSCs between FSC layers. (A-F) Flies with *UAS-H2B-RFP*, *actin-GAL4* and *tub-tsGAL80* were kept at 29C for 4d and then moved to 18C for the times indicated. Ovaries were stained for Traffic Jam (not shown) and Fas3 (blue). (A) Prior to 18C chase, H2B-RFP showed strong expression in layer 2 and 3 FSCs (blue and yellow arrows) and region 2a ECs, weaker expression in layer 1 FSCs (yellow arrow) and even weaker expression in FCs, which express Fas3 strongly. (B, C) Germaria from flies kept at 18C for 7d had no detectable H2B-RFP in FCs, and variable dilution of H2B-RFP across FSC layers, ranging from minimal for the FSCs indicated by arrows in B, to strong dilution of FSCs in layer 1 (white arrows in C) and layer 2 (bottom blue arrow in C). (D) Germarium from a fly kept for 12d at 18C showing large differences between FSCs in a single layer. Exceptional FSCs have high RFP expression in layer 1 (top white arrow) and weak RFP expression in layer 3 (middle yellow arrow). Layer 2 FSCs (blue arrows) had a mixture of strong and weak expression (blue arrows), and most layer 1 cells had very weak expression (bottom white arrow). (E) Germarium kept at 18C for 12d showing diluted layer 1 and 2 FSCs (top white and blue arrows) and a layer 1 FSC (bottom white arrow). (F) Germarium kept for 21d at 18C showing strongly reduced H2B-RFP in layers 1-3 FSCs (top) and layers 1-2 (bottom) but one strong layer 3 cell (bottom). (G) Live imaging frames at 0,4 and 8h of a germarium kept for 21d at 18C shows a layer 1 cell (white arrow) moving into layer 2, and a layer 2 cell (blue arrow) moving into layer 1. Layers were determined by H2B-RFP expression level, with most layer 1 cells expressing very weak RFP and layers 2 and 3 expressing stronger RFP. (H) Live imaging of a germarium kept at 18C for 16d shows a layer 2 FSC (white arrow) moving to layer 3 at 1h 31min and to region 2a at 3h 11min. A layer 3 FSC is indicated by the yellow arrow for reference. All scale bars, 20μm.

H2B-RFP signal was largely cleared from early FCs to undetectable levels within 7d of chase incubation at 18C (Fig. 5B). ECs and anterior FSCs retained a strong H2B-RFP signal, while intensity in some layer 1 FSCs was markedly lower but clearly detectable (Fig. 5B, C). That is consistent with an estimated cell cycle time of under 11h for FCs (47), compared to our estimate of nearly 16h for layer 1 FSCs and over 50h for more anterior FSCs. The intensity of H2B-RFP staining was measured in layer 1 and 2 FSCs, normalizing to layer 3 intensity in overlapping z-sections. Prior to chase, average relative H2B-RFP intensities were 1 (layer 3), 0.91 (layer 2), 0.69 (layer 1) and 0.41 (early FCs). After 7d at 18C, layer 1 and 2 values had dropped on average to 0.33 and 0.56, respectively (relative to layer 3 in the same samples), equivalent to 48% (layer 1) and 61% (layer 2) of pre-chase values and indicating progressively faster division from layer 3 to layer 1. By eye, H2B-RFP signal was much weaker in some layer 1 FSCs than others. This visual threshold corresponded to a measured dilution of at least four-fold from the pre-chase average, with 45% (5/11) of layer 1 FSCs below the threshold. Using the same criterion of at least four-fold dilution, only 15% (2/13) of layer 2 and no (0/10) layer 3 cells were below the threshold.

By 12d at 18C, the average H2B-RFP signal, normalized to strong layer 3 cells and expressed relative to pre-chase levels, had declined to 25% in layer 1 and 51% in layer 2. The proportion of cells with greatly diluted signal (more than 4-fold) was 61% (8/13) in layer 1, 17% (2/12) in layer 2 and none (0/14) in layer 3 (Fig. 5D, E). Clearly, whether measuring average intensity of all cells in a layer or the proportion with greatly reduced H2B-RFP signal, the results show a gradient of division from layer 1 (high) through layer 2 (intermediate) to layer 3 (low).

By 21d at 18C, even some layer 3 cells had very reduced H2B-RFP signal and each layer showed prominent mosaicism (Fig. 5F). We therefore scored the proportions of cells in each layer that appeared visually to have very diluted signal. Greatly diluted signal was observed for 18% (4/22), 25% (9/36) and 53% (17/32) of layer 3 cells at 25d, 32d and 34d. H2B-RFP-depleted layer 2 cells were more frequent at 74% (14/19), 100% (15/16) and 74% (17/23), while almost all layer 1 cells had this property (88% (30/34), 100% (31/31) and 100% (39/39) at the three time-points.

Particularly at late times, there were prominent differences in H2B-RFP signal among cells in the same AP layer. This could result from non-uniform division rates, possibly stochastic in origin, non-uniform initial labeling intensity or movement of cells between layers. Previously, movement between FSC layers was inferred from observing that two derivatives of a single labeled FSC were observed in different layers 3d after marking at a frequency of about 50% (10). Also, in multiple MARCM lineage analyses, including lineages expressing an apoptosis inhibitor, the rate of production of marked ECs per anterior marked FSC appeared to reach a steady state by about 6d, suggesting that FSC-derived ECs also revert to FSC status (18). Here, we used samples, which also included a *ubi-GFP* marker to visualize all somatic cells, and had been chased for at least 7d at 18C to see if we could directly observe AP cell movements by live imaging. The pattern of H2B-RFP staining was helpful to define the location of layer 1 and other layers, and also provided evidence of a tracked cell having the initial assigned identity by virtue of its inferred division history. We observed instances of a layer 2 FSC moving into layer 1, and vice versa (Fig. 5G). Movement in both directions is consistent with our inference that the division and differentiation rates are balanced within each layer, and the consequent expectation of no net AP flow. We also observed a layer 2 FSC moving from layer 2 into layer 3 and then into the location of a region 2a EC (Fig. 5H) and a layer 3 cell moving into layer1 (not shown).

### MARCM clonal analysis of responses to regulators of FSC cycling

The mechanisms that regulate FSC proliferation can also be probed by clonal analyses, using the MARCM technique (48), where the behavior of GFP-marked FSCs of altered genotype is measured in the context of unmarked normal cells (Fig. 6B, C). We have undertaken such assays here and in the past, using a standard protocol with measurements at 6d and 12d after clone induction (18). Division rate is indicated by the EdU index of marked cells in specific locations at 6d. Our live analyses of FUCCI reporters suggests EdU index can be a good qualitative indicator of changes in division rate within a given layer, but may be quantitatively inaccurate because of changes to the length of S-phase. In clonal analyses, we also measure relative rates of conversion of FSCs to ECs and to FCs (at 6d), changes in the AP distribution of genetically altered marked FSCs (at 12d), and whether the marked FSC population grows or declines in competition with unmarked FSCs (by 12d) (18). These additional measurements (Fig. 6A) allow appraisal of whether factors affecting FSC division rate also affect other FSC behaviors, and whether the net effect on the number of marked FSCs suggests an accord between FSC proliferation and the division rate indicated by EdU index.

**Figure 6.**
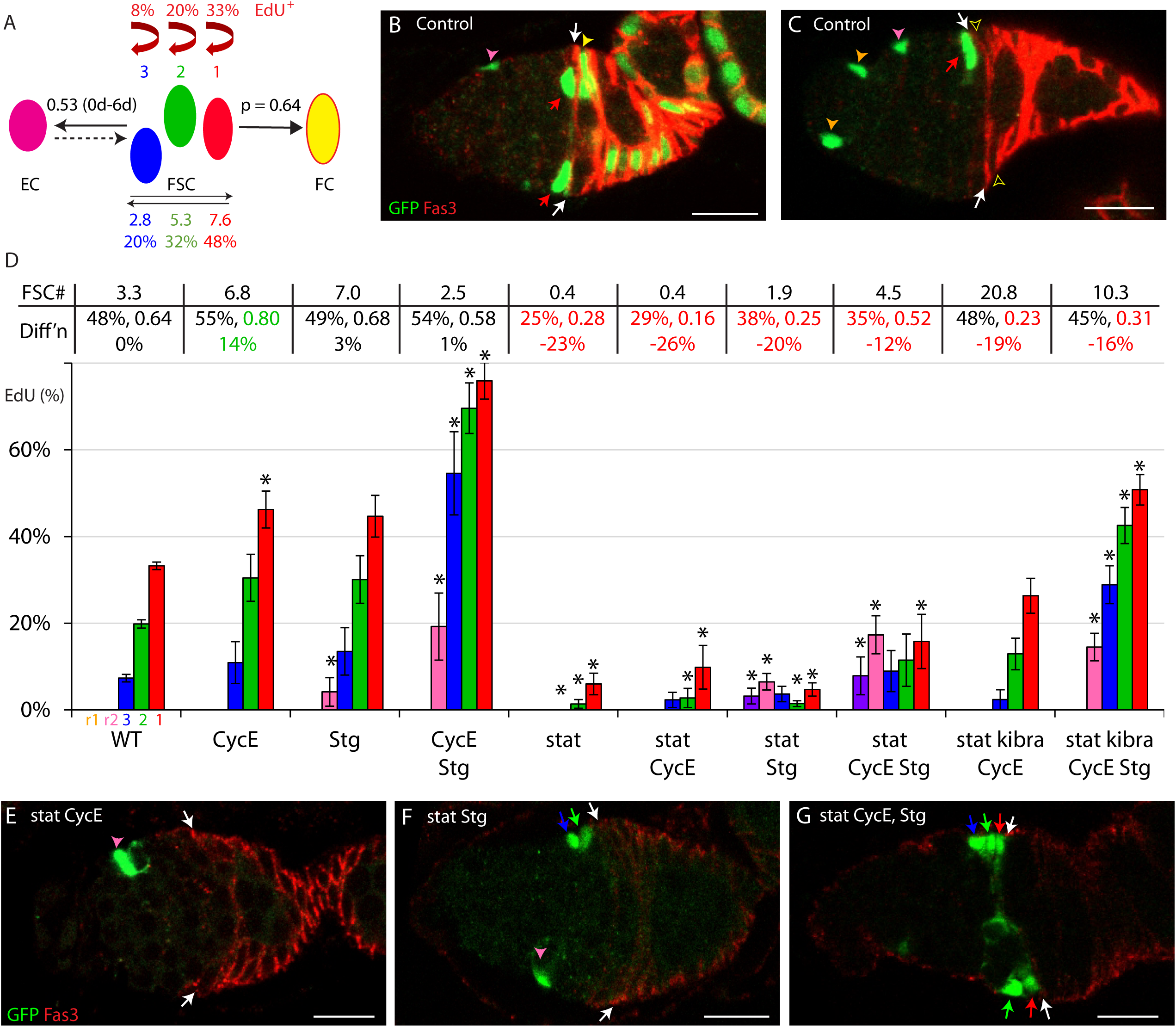
Restoration of normal division rates to *stat* mutant FSCs requires both excess G1/S and G2/M regulators. (A) Illustration of separately measured parameters of FSC behavior from multiple control MARCM experiments reported here and previously (18); EdU indices in each FSC layer, average number and percentage of all FSCs in each layer, ECs produced per anterior FSC from 0-6d, and inferred probability (p) of conversion of a single layer 1 FSC to an FC in each 12h budding cycle. (B, C) Control MARCM samples, illustrating GFP-marked (green) layer 1 FSCs (red arrows), immediate FCs (filled yellow arrowhead in B, none in C; empty arrowhead) just posterior to the anterior border (white arrows) of strong Fas3 (red) staining, r2a (pink arrowheads) and r1 ECs (orange arrowheads, only in C). (D) Percentage of marked cells of the indicated genotypes that incorporated EdU for cells in the location of r1 ECs (purple), r2a ECs (pink), layer 3 (blue), layer 2 (green) and layer 1 (red) FSCs. SEMs and significant differences from control values are indicated (*, p<0.05, n-1 chi-squared test). For each genotype, above the graph, the average number of marked FSCs at 12d is indicated, as well as the percentage of FSCs in layer 1 at 6d, the calculated probability of FSC to FC conversion per cycle and (below those numbers) the inferred percentage loss of FSCs due to altered FC differentiation per 12h budding cycle. Large increases (green) or decreases (red) are highlighted. (E-G) MARCM samples for *stat* mutant FSCs expressing excess (E) CycE, (F) Stg or (G) both, with the Fas3 (red) border indicated (white arrows). (E) Loss of STAT activity caused greatly reduced division and anterior displacement of marked cells. (G) Excess CycE and Stg together partly restored FSC numbers (red arrow-layer 1; green arrow-layer 2; blue arrow-layer 3), with (F) much less restoration by excess Stg alone. Scale bars 10μm.

For example, in previous studies it was found that increased CycE raised the EdU index, did not substantially affect conversion of FSCs to ECs or FCs, and resulted in amplification of the marked FSC population (18). By contrast, loss of Wnt signaling did not significantly affect the FSC Edu index but increased conversion of layer 1 FSCs to FCs and caused most FSCs to move posteriorly into layer 1, resulting in a net decline of marked FSCs (10, 18). The effects of altered JAK-STAT signaling were more complex. Loss of STAT drastically reduced the EdU index but loss of marked FSCs was tempered by reduced conversion of FSCs to FCs. Conversely, the increased EdU index in both FSCs and ECs caused by increased JAK-STAT signaling increased was partially countered by increased conversion of FSCs to FCs, but still led to a large increase in marked FSCs (18).

Here we examined the properties of FSC clones in response to a number of potential regulators of FSC division rate in order to understand better how major signals regulate FSC division, and whether other FSC properties are simultaneously affected.

### JAK-STAT stimulation of EdU index depends on G1/S and G2/M actions

Increased CycE (EdU index of layer1, layer2, layer3 marked FSCs= 46%, 30%, 11%) and increased Stg (45%, 30%, 13%) each elevated the EdU index of FSCs in all layers (controls: 33%, 20%, 7%) but together they produced an even larger increase (76%, 70%, 55%), higher than observed with any other experimental manipulation (Fig. 6D). In these MARCM clonal tests, transgenes are driven by a combination of *tub-GAL4* and *act-GAL4*, and are likely expressed at similar levels among all FSC layers and ECs. The tests described earlier with FUCCI reporters used *C587-GAL4*, which is expressed at higher levels in anterior FSCs and ECs than in posterior FSCs. Consequently, the effects on EdU index were more pronounced in posterior cells in clonal analyses. Nevertheless, the effects on EdU index were broadly similar for excess CycE and Stg, with the common conclusion that stimulating both G1/S and G2/M transitions has an additive effect on promoting FSC cycling. Among ECs, there was no response to excess CycE (Fig. 6D). Excess Stg triggered some S-phase entry for r2a ECs but addition of excess CycE further increased the EdU index (from 4% to 19%). The most anterior, r1 ECs were not found in S phase even with excess CycE and Stg.

We then tested whether deficiencies in JAK-STAT signaling could be compensated by excess CycE or Stg. As described previously (18), excess CycE only very weakly increased the EdU index for *stat* mutant FSCs (9.8%, 2.8%, 2.3% vs 6.0%, 1.4%, 0% for layers 1, 2 and 3, respectively) (Fig. 6D, E). Excess Stg produced a similar result (4.7%, 1.5%, 3.7%) (Fig. 6D, F). However, EdU incorporation was significantly increased for *stat* mutant FSCs, especially in more anterior locations, by providing excess CycE and Stg together (15.8%, 11.5%, 9.0%) (Fig. 6D, G).

This synergy was tested further in the background of an additional genetic alteration. Yki activation through the Hippo pathway (using a *kibra* mutation) was described previously to enhance EdU incorporation in *stat* mutant FSCs alone (6.5%, 4.7%, 0.7%), and more potently together with excess CycE (26.3%, 12.9%, 2.4%) (18). There is some evidence that Yki acts in part in FSCs through increasing transcriptional induction of *cycE* (17). Addition of both *UAS-Stg* and *UAS-CycE* to *kibra stat* mutant FSCs increased EdU incorporation to levels significantly higher than normal (51%, 43%, 29% vs 33%, 20%, 7%) (Fig. 6D). Thus, the normal input from JAK-STAT signaling was compensated by stimulation of both G1/S and G2/M transitions but only poorly by reagents expected to act on only one phase transition. Notably, even though the normally graded input from posterior (high) to anterior (low) of JAK-STAT was replaced by excess CycE and Stg driven by a constitutive promoter (together with uniform Hippo pathway inactivation when *kibra* was altered), the EdU index of posterior FSCs remained higher than that of anterior FSCs in all cases, indicating strong graded influences other than JAK-STAT.

EdU incorporation into r2a ECs was not stimulated by excess Stg when *stat* activity was absent. However, *stat* mutant ECs did incorporate EdU when both *UAS-Stg* and *UAS-CycE* were provided in the presence (10%) or absence of a *kibra* mutation (14%), including some r1 ECs (8%) in the latter case (Fig. 6D). Thus, EC cycling can be induced by promoting both G1/S and G2/M transitions. The declining levels of JAK-STAT pathway activity are normally insufficient in ECs but can supplement responses to excess Stg, suggesting a modest contribution to G1/S transitions in region 2a ECs.

If a specific genetic manipulation has no effect on FSC location or conversion to ECs or FCs, we would expect the change in marked FSC number over time to depend only on FSC division rate relative to wild-type FSCs, which is indicated, at least qualitatively by the EdU index. Layer 1 FSCs should have the most impact because they normally divide more than three times faster than anterior FSCs. The default correlation between EdU index and marked FSC number can be disrupted most potently by changes in differentiation of FSCs to FCs because FCs are normally produced at about four times the frequency of ECs (4). We therefore report in Fig. 6D and similar later graphs, the percentage of FSCs in layer 1 (normally 48%, Fig. 6A) and the probability of a single layer 1 FSC becoming a FC in one cycle of egg chamber budding (p=0.64 normally, Fig. 6A) as measures of FC production (“Diff’n”) (18). Below those values is the calculated expected percentage change in FSC loss due to FC formation relative to wild-type per cycle of egg chamber budding (a positive number indicates greater loss; see Methods). In the absence of functional STAT, there is reduced FC differentiation (18). That property was substantially retained with excess CycE, Stg, or both (Fig. 6D). Consequently, we would expect that the number of marked FSCs correlates with the EdU index for all of these *stat* mutant genotypes if EdU indices are reporting division rates reasonably accurately. Indeed, provision of excess CycE and Stg together increased *stat* mutant FSC numbers by more than either alone, supporting the inference from EdU indices that FSC proliferation is enhanced by complementing both a G1/S and G2/M transition defect due to the absence of functional STAT.

In the presence of normal STAT activity, excess CycE and excess Stg each increased the EdU index and the number of marked FSCs. However, in the presence of both factors and an extremely high EdU index, with no significant change to FC differentiation, there was no increase in FSC numbers (Fig. 6D). This artificial manipulation aimed at greatly speeding G1/S and G2/M transitions, in contrast to manipulating natural signals (see below), appears therefore to produce an unproductive imbalance that ultimately limits FSC proliferation, perhaps due to cell death or greatly prolonged S phase.

### Preferential effects of Yki on anterior FSC cycling

The Hippo pathway includes a variety of sensors of cell crowding, tension, polarity and other mechanical signals, including Kibra, which promote activation of Hippo (Hpo) and Warts (Wts) protein kinases, leading to Yki phosphorylation and restriction of Yki nuclear access as a common central response (29). It was previously shown that inactivation of Yki led to quiescence and apoptosis of FSCs, whereas inactivation of *kibra*, *wts* or *hpo* increased FSC proliferation (17). However, those studies preceded re-evaluation of FSC numbers, locations and behavior and therefore assayed only a fraction of all FSCs (10). Here we found that *yki* mutant FSCs had greatly reduced EdU incorporation in all layers (8%, 0%, 0% vs 33%, 20%, 7%) and were reduced in number (0.7 vs 3.3), without large changes in FSC proportion in layer 1 (42% vs 48%), or differentiation frequency to FCs (0.50 vs 0.62) or to ECs (0.67 vs 0.55) (Fig. 7A). By contrast, FSC numbers were increased to a similar degree by loss of *kibra* (7.3), wts (7.7) or *hpo* (5.0) (Fig. 7A, C, D) without greatly affecting FSC proportion in layer 1 (average 55%), or the differentiation frequency to FCs (average 0.58) or ECs (average 0.70). These results are in accord with earlier conclusions (17). However, the stimulation of the EdU index by *kibra*, *wts* and *hpo* was not as large as reported previously. A weighted average among all three mutant genotypes showed only a very small increase in layer 1 (from 33% to 35%; n=541) and layer 2 EdU indices (from 20% to 22%; n=345), with a somewhat greater impact in layer 3 (7% to 14%; n=167). These results suggest that the restraint of Yki activity by the Hippo pathway may normally be highest or most effective in limiting FSC division in layer 3 FSCs. It is possible that increased division of anterior FSCs is sufficient for the observed FSC amplification. However, deductions from live imaging of FUCCI reporters showed that the EdU index is not necessarily a reliable quantitative measure of FSC division rate. It is therefore also possible that reduction of Hippo pathway activity promotes FSC division more than reflected by measuring the EdU index. No EdU incorporation was seen for *kibra*, *hpo* or *wts* mutant ECs.

**Figure 7.**
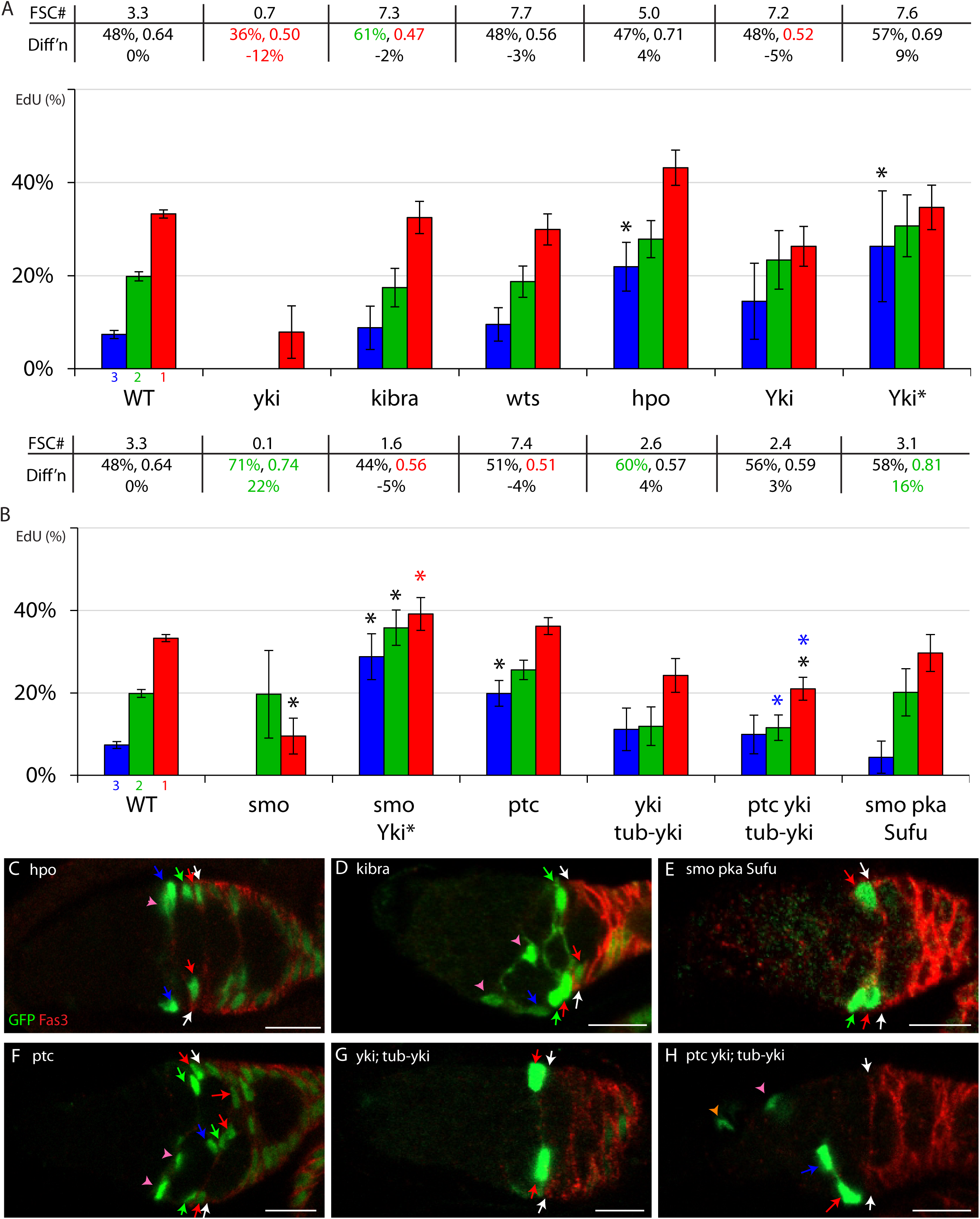
Excess Hh signaling and Yki activity modestly increase anterior FSC EdU indices. (A, B) Percentage of marked cells of the indicated genotypes (Yki indicates *UAS-Yki*; Yki* indicates *UAS-YkiS168A*) that incorporated EdU for cells in the location of r1 ECs (purple), r2a ECs (pink), layer 3 (blue), layer 2 (green) and layer 1 (red) FSCs. SEMs and significant differences from control values for cells in a given location are indicated (*, p<0.05, n-1 chi-squared test). Additional pair-wise comparisons show significant responses to a signal alteration in an altered candidate mediator background (red asterisk; *ptc yki; tub-yki* vs *yki; tub-yki*, *smo; UAS-Yki\** vs *smo*) and whether the response is significantly lower than in a wild-type background (blue asterisks; *ptc yki; tub-yki* vs *ptc, smo; UAS-Yki\** vs *UAS-Yki\**). Absence of colored asterisks indicates no significant differences. For each genotype, above the graph, the average number of marked FSCs at 12d is indicated, as well as the percentage of FSCs in layer 1 at 6d, the calculated probability of FSC to FC conversion per cycle and (below those numbers) the inferred percentage loss of FSCs due to altered FC differentiation per 12h budding cycle. Large increases (green) or decreases (red) are highlighted. (C-H) MARCM samples for the indicated genotypes, with the Fas3 (red) border indicated (white arrows) and colored arrows indicating layer 1 (red), 2 (green) or 3 (blue) FSCs, r2a (pink arrowheads) and r1 (orange arrowheads) ECs. FSC numbers were (C, D) increased by *hpo* and *kibra* mutations, (E) similar to control values for *smo pka Su(fu)* FSCs, which have roughly normal pathway levels but do not respond to Hh, and (F) increased by loss of *ptc*. Replacement of normal *yki* gene activity with *tub-yki*, eliminating normal transcriptional control, (G) did not greatly alter FSC numbers but (H) suppressed the increase normally elicited by increased Hh signaling (loss of *ptc*). Scale bars 10μm.

Increasing Yki levels with *UAS-Yki* also increased FSC numbers (to 7.2) and increased the layer 3 FSC index (15% vs 7%), with little or no increase in the EdU index of layer 1 (26% vs 33%) or 2 (23% vs 20%), similar to the effects of Hippo pathway mutations (Fig. 7A). Expression of *UAS-Yki-S168A*, which is resistant to a major form of inactivation by the Hippo pathway (29), increases both Yki levels and by-passes restraint by the Hippo pathway. This also increased FSC numbers (to 7.6) and produced a larger increase in EdU incorporation that was again more prominent in layers 3 (26% vs 7%) and 2 (31% vs 20%) than in layer 1 (35% vs 33%) (Fig. 7A). Neither excess Yki nor activated Yki induced EdU incorporation in ECs. The whole set of results (Fig. 7A) supports the earlier conclusions that Yki is required for significant FSC cycling and maintenance, that increasing Yki activity through additional *yki* transcription or relief from Hippo pathway inhibition, or both, significantly increased FSC numbers without much effect on FSC location or differentiation to ECs or FCs. The observed increase in EdU incorporation for genotypes with increased Yki activity was smaller than previously found by examining only a subset of FSCs (17) and was mostly evident only in the most anterior FSCs.

### Hedgehog Signaling acts through Yki in FSCs but graded activity is dispensable

It was previously shown that Hedgehog (Hh) signaling normally provides strong stimulation of FSC proliferation and does so principally by inducing *yki* transcription (16, 17, 19, 26). Here we present a more detailed analysis, benefiting from the revised, current picture of FSC organization (10). Loss of Hh signaling in *smo* mutant FSC clones led to severely reduced FSC numbers (0.1 vs 3.3) and EdU incorporation (Fig. 7B). Addition of activated Yki (*UAS-YkiS168A*) restored EdU incorporation to levels greater than controls in all FSC layers (39%, 36%, 29%) and restored FSC numbers (2.6) almost to normal, with a roughly normal proportion of FSCs in layer 1 (42%) and normal differentiation frequencies to FCs (0.56) and ECs (0.90).

Full Hh pathway activation in *ptc* mutant clones produced responses very similar to Hippo pathway inactivation, with only a small increase in EdU incorporation overall, but most prominently in layer 3 (20% vs 7%) compared to layers 1 (36% vs 33%) and 2 (26% vs 20%), a significant increase in FSC number (7.4 vs 3.3) and no major changes to the proportion of FSCs in layer 1 (51%) or differentiation to FCs (0.51) or ECs (0.46) (Fig. 7B, F). *ptc* mutant ECs did not incorporate EdU.

A *tub-yki* transgene rescues *yki* null animals to produce viable and fertile flies (49). *yki* mutant FSC clones in animals carrying the *tub-yki* transgene maintained an almost normal number of FSCs (2.7 vs 3.3), proportion of FSCs in layer 1 (50%), FC (0.58) and EC (0.68) differentiation frequencies (Fig. 7B, G). EdU incorporation was lower than controls in layers 1 (24% vs 33%) and 2 (12% vs 20%) but slightly higher in layer 3 (11% vs 7%). In this background of *yki* activity provided to marked FSCs only by the *tub-yki* transgene, the loss of *ptc* activity no longer increased FSC number (2.4 vs 2.7) or EdU incorporation in layer 1 (21% vs 24%), 2 (12% vs 12%) or 3 FSCs (10% vs 11%) (Fig. 7B, H). These results support the previous conclusion (17) that increased Hh signaling elicits greater FSC cycling and consequently increased FSC numbers through transcriptional induction of *yki* and that normal Hh pathway activity likely acts substantially in the same manner.

We also tested a genotype, *smo pka; Su(fu)*, which was previously deduced to reproduce roughly normal levels of Hh pathway activity in FSCs (26), but without any possibility for modulation by Hh because of the absence of functional Smo. We found a roughly normal pattern of EdU incorporation among FSC layers (30%, 20%, 4%), albeit with slightly reduced values, and roughly normal FSC numbers (3.1 vs 3.3), indicating that the shallow gradient of Hh signaling (declining from anterior to posterior) is not of major importance to supporting differential cycling among FSC layers (Fig. 7B, E). *smo pka* FSC clones, also unable to sense Hh levels, but with lower overall Hh pathway activity because the inhibitor Su(fu) is present (26), had substantially lower EdU incorporation in all layers (15%, 10%, 6%) and reduced FSC numbers (0.5) (Fig. 9B), confirming previous conclusions that the magnitude of Hh signaling is nonetheless critical to maintain normal rates of FSC cycling (26).

### PI3K pathway stimulation of FSC cycling independent of CycE or Yki induction

Both reduced and increased PI3 kinase pathway activity have been reported to affect FSC division and survival (20). Here, we examined those consequences in more detail. Chico acts downstream of the insulin receptor to stimulate PI3K pathway activity (42, 45). Loss of *chico* activity severely reduced EdU incorporation in all FSC layers (18%, 4%, 0%) (Fig. 8A). FSC numbers were also reduced (1.0 vs 3.3), but may be bolstered by significantly reduced conversion of layer 1 FSCs to FCs (0.35 vs 0.64). Addition of excess CycE did not increase EdU incorporation (22%, 9%, 0%), FSC numbers (1.2) or FC differentiation frequency (0.33) significantly, suggesting that PI3K pathway deficiency produces a cycling defect that is not remedied by excess CycE (Fig. 8A). That result is consistent with the inference from FUCCI reporter studies that PI3K activity appears principally to regulate the G2/M transition.

**Figure 8.**
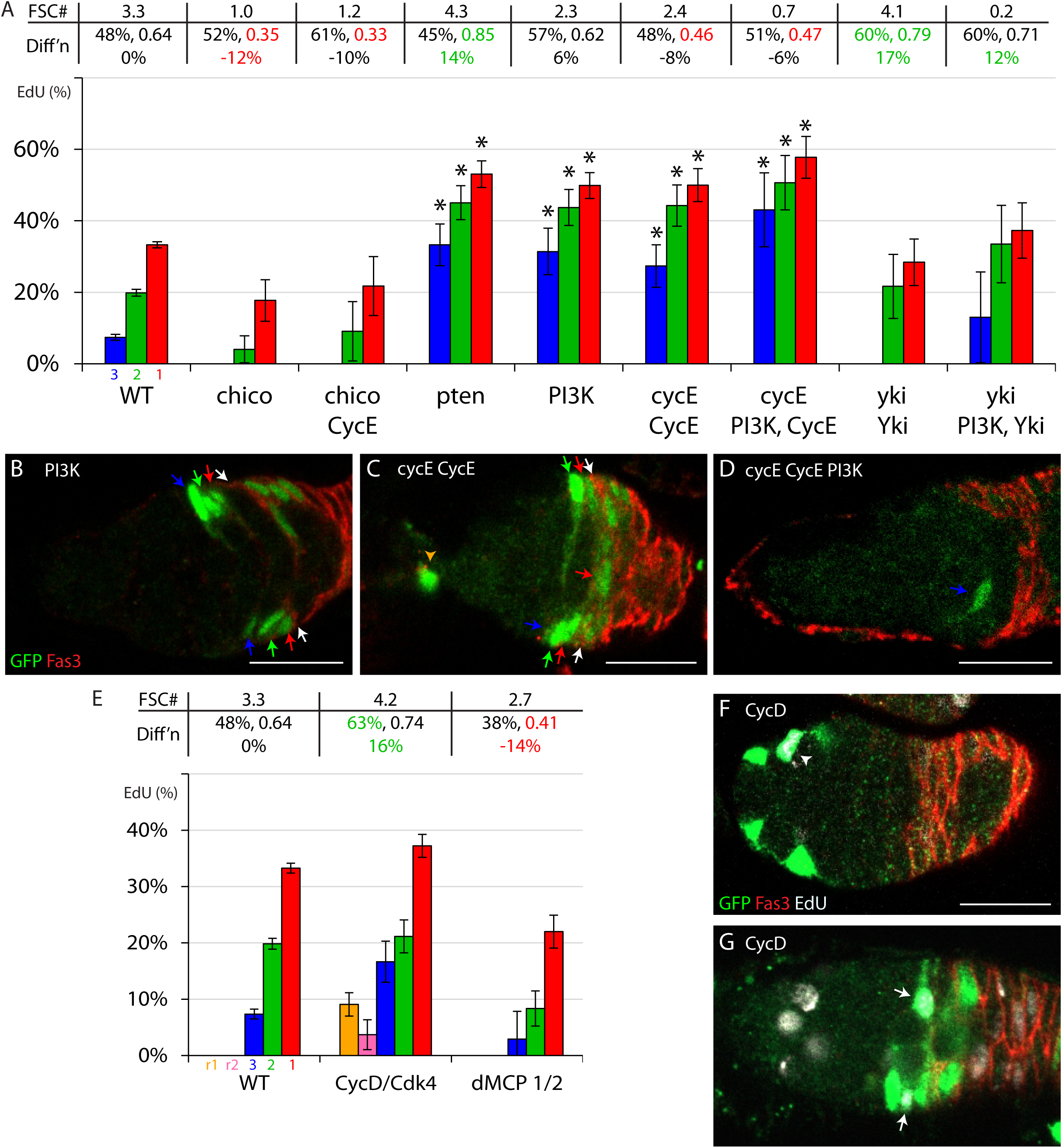
PI3K pathway stimulation of FSC EdU incorporation does not require transcriptional regulation of *cycE* or *yki*. (A, E) Percentage of marked cells of the indicated genotypes (CycE, PI3K, Yki, CycD, Cdk4 all indicate the named *UAS*-driven transgenes) that incorporated EdU for cells in the location of r1 ECs (purple), r2a ECs (pink), layer 3 (blue), layer 2 (green) and layer 1 (red) FSCs. SEMs and significant differences from control values are indicated (*, p<0.05, n-1 chi-squared test). Additional pair-wise comparisons show significant responses to a signal alteration in an altered candidate mediator background (red asterisk; *cycE; UAS-PI3K UAS-CycE* vs *cycE; UAS-CycE*, *yki; UAS-PI3K UAS-Yki* vs *yki; UAS-Yki*, *chico; UAS-CycE* vs *chico*) and whether the response is significantly lower than in a wild-type background (blue asterisks; *cycE; UAS-PI3K UAS-CycE* vs *UAS-PI3K*; *yki; UAS-PI3K UAS-Yki* vs *UAS-PI3K*). Absence of colored asterisks indicates no significant differences. For each genotype, above the graph, the average number of marked FSCs at 12d is indicated, as well as the percentage of FSCs in layer 1 at 6d, the calculated probability of FSC to FC conversion per cycle and (below those numbers) the inferred percentage loss of FSCs due to altered FC differentiation per 12h budding cycle. Large increases (green) or decreases (red) are highlighted. (B-D, F, G) MARCM samples for the indicated genotypes, with the Fas3 (red) border indicated (white arrows) and (B-D) colored arrows indicating layer 1 (red), 2 (green) or 3 (blue) FSCs, r2a (pink arrowheads) and r1 (orange arrowheads) ECs. FSC numbers were (B) increased in some samples by excess PI3K, but (A) not on average. (C) FSC numbers were increased by replacing *cycE* function with *UAS-CycE*, eliminating normal transcriptional control. Under these conditions, (D) excess PI3K greatly reduced FSC numbers. (F, H) Samples also labeled with EdU (white) show (F) significant EdU labeling of ECs (white arrowhead) and (G) more layer 3 FSC (white arrows) labeling in response to excess CycD and Cdk4. Scale bars 10μm.

Activation of the PI3K pathway through loss of *pten* function (values are averages for strong alleles *dj* and *c494*) increased EdU incorporation in all layers (53%, 45%, 33%). It also increased FC differentiation frequency (0.85 vs 0.64), with unchanged EC differentiation (0.56) and FSC proportion in layer 1 (45%). FSC numbers were increased modestly (3.3 to 4.3), consistent with increased FC differentiation partially offsetting the effects of increased FSC division rate (Fig. 8A). These results are precisely the converse of the consequences of reducing PI3K activity. It is interesting that loss of Pten activity, like JAK-STAT activity promotes both FSC cycling and differentiation to FCs.

The consequences of expressing excess PI3K p110 catalytic subunit (*UAS-PI3K*) were similar to *pten* loss for the FSC EdU index (50%, 44%, 31%). However, there was no increase in FC differentiation as seen for *pten,* and FSC numbers declined slightly (2.3 vs 3.3) (Fig. 8A, B). The pattern and magnitude of pathway changes induced by loss of Pten and increased PI3K catalytic subunit are not expected to be identical, but we have no obvious explanation of the failure of FSCs to accumulate in response to excess PI3K.

The response to *UAS-PI3K* was examined in a *cycE; UAS-CycE* background to test whether transcriptional induction of *cycE* was important. The EdU index of *cycE; UAS-CycE* (50%, 44%, 27%) was already higher than normal but was increased by the addition of *UAS-PI3K* (58%, 51%, 43%), suggesting that PI3K can affect FSC cycling without altering *cycE* transcription (Fig. 8A, C, D). The increase of EdU indices due to excess PI3K was, however, not statistically significant. PI3K activity has been found to promote Yki activity in FCs and wing discs (50). A similar experiment was therefore performed to test regulation through *yki* transcription. The EdU index of *yki; UAS-Yki* (28%, 22%, 0%) FSCs was lower than normal and was increased by the addition of *UAS-PI3K* (37%, 33%, 13%), suggesting that PI3K can also affect FSC cycling without altering *yki* transcription (Fig. 8A). The EdU indices for *yki; UAS-Yki + UAS-PI3K* were intermediate between those of *yki; UAS-Yki* and *UAS-PI3K* alone, and neither difference reached statistical significance in these tests. It is believed that Yki acts substantially by inducing *cycE* transcription in FSCs (17), so the results of tests that prevent normal transcriptional regulation of *cycE* and *yki* both indicate, albeit short of clear statistical significance, actions of PI3K independent of *cycE* induction. This conclusion is consistent with deductions from FUCCI reporters that PI3K pathway activity appears principally to stimulate the G2/M transition.

In both *cycE; UAS-CycE* and *yki; UAS-Yki* backgrounds, the addition of excess PI3K reduced the number of marked FSCs despite increasing the FSC EdU index (Fig. 8A-D). There was no evident increase in FC differentiation in either case. So, the unexpected reduction in FSC numbers for a given EdU index, which was also observed in response to excess PI3K in a normal genetic background, is not readily explained by measured parameters of FSC behavior (which do not include cell death).

### Growth and Metabolism

Cell proliferation requires growth and accumulation of suitable macromolecules, supported by appropriate metabolic activity, in addition to cell cycling. It is likely that some signals received by FSCs stimulate growth and alter their metabolism in parallel to effects on cell cycling, and possible that growth or metabolic responses also connect to cell cycling. Indeed, it is possible that the failure of excess CycE together with String to stimulate FSC accumulation despite very high rates of EdU incorporation is because of a failure to stimulate growth. By contrast, the signaling pathways regulating FSC cell cycles may additionally stimulate growth; the PI3K pathway has especially strong connections to ribosome assembly and protein synthesis rates in other settings (42).

In Drosophila wing discs, CyclinD/Cdk4 stimulates cycling (involving Rb phosphorylation) and growth, primarily through stimulating mitochondrial activity (51–53). We found that increased expression of CycD together with Cdk4 modestly increased FSC EdU incorporation (to 37%, 21%, 17%) and FSC accumulation (to 4.2). Given this modest stimulation, it was surprising that EdU incorporation was present in ECs (to 6%) (Fig. 8E-G). This response suggests that excess CycD/Cdk4 promotes the G2/M transition, like the only other known effectors of EC cycling (JAK-STAT or Stg). Some of the EdU positive ECs were in region 1 (as seen for JAK-STAT but not for Stg), where all cells are normally in G1, suggesting that the G1/S transition may also be stimulated.

The mitochondrial pyruvate carrier (MPC) can import pyruvate to promote mitochondrial oxidative phosphorylation (54, 55). In Drosophila intestinal stem cells, loss of the dMCP1 subunit of the heterodimeric carrier protein increased proliferation. We found that the same dMCP1 loss of function mutation in FSCs decreased EdU incorporation (to 22%, 8%, 3%), reduced FC differentiation and modestly reduced FSC accumulation (to 2.6), suggesting that oxidative phosphorylation contributes to promote normal FSC cycling and FSC differentiation to FCs (Fig. 8E). Prior studies showed that many components contributing to oxidative phosphorylation are important for FSC survival but the underlying reasons are likely varied, including in some cases detrimental Reactive Oxygen Species (ROS) production, and incompletely investigated (20). Although the regulation and responses to mitochondrial function in FSCs promise to be complex, the results from eliminating MCP activity provide a strong indication that FSCs do not rely on glycolysis alone, and it remains to be determined whether the stimulatory effect of CycD/Cdk4 activity on FSC cycling may be mediated by mitochondrial responses, as in some other settings.

### JAK-STAT stimulates FSC cycling partly through *yki* regulation

Increased expression of JAK (*UAS-Hop*) has previously been shown to increase STAT activity and promote increased FSC division as well as initiate division in EC territory (18). Here we explored whether these effects rely on transcriptional induction of *yki*, CycE/Cdk2 activity, Hh pathway activity or integrin signaling. In *yki; tub-yki* FSCs, where *yki* is not subject to normal transcriptional regulation, addition of *UAS-Hop* increased the FSC index in each layer (29%, 17%, 14% vs 24%, 12%, 11%) but to much lower levels than in FSCs expressing *UAS-Hop* alone (50%, 40%, 32%) (Fig. 9A, C, F). FC differentiation frequency was greatly increased (1.0 vs 0.57), as for *UAS-Hop* alone, but FSC numbers were unchanged (2.7), contrasting with *UAS-Hop* alone (11.2 FSCs). Thus, transcriptional induction of *yki* appears to be a significant component of the response to increased JAK-STAT pathway activity. A similar test was made using *UAS-Yki* instead of *tub-yki*. The greater EdU indices and FSC numbers observed for *yki* complementation by *UAS-Yki* compared to *tub-yki* suggest that *UAS-yki* produces more *yki* gene product in FSCs than either *tub-yki* or the normal *yki* gene (Fig. 9A, H). The addition of *UAS-Hop* was now more effective at increasing EdU incorporation of *yki; UAS-yki* FSCs (from 32%, 18%, 0% to 41%, 29%, 20%) and FSC numbers (from 5.2 to 10.9), while *UAS-Hop* still increased FC differentiation frequency markedly (from 0.71 to 0.94) (Fig. 9A, G). Thus, JAK-STAT can also strongly stimulate FSC cycling independent of *yki* induction, and does so more effectively when Yki levels are higher. However, the change in EdU index in FSCs was still reduced in the absence of the potential to induce *yki* transcription even under these conditions of an elevated baseline of *yki* expression. These results suggest that stimulation of FSC division by additional JAK-STAT signaling depends partly on *yki* induction. Provision of excess Yki did not increase the very low EdU indices of *stat* mutant FSCs (4.3% vs 6.0% in layer 1) but did substantially increase the average number of labeled FSCs at 12d after clone induction (from 0.4 to 2.5) (data not shown). Addition of *tub-yki* increased *stat* mutant FSC number to only 1.2 and also failed to increase the EdU index (data not shown). Clearly, mediators other than increased Yki are important to the normal proliferative input of JAK-STAT signaling.

**Figure 9.**
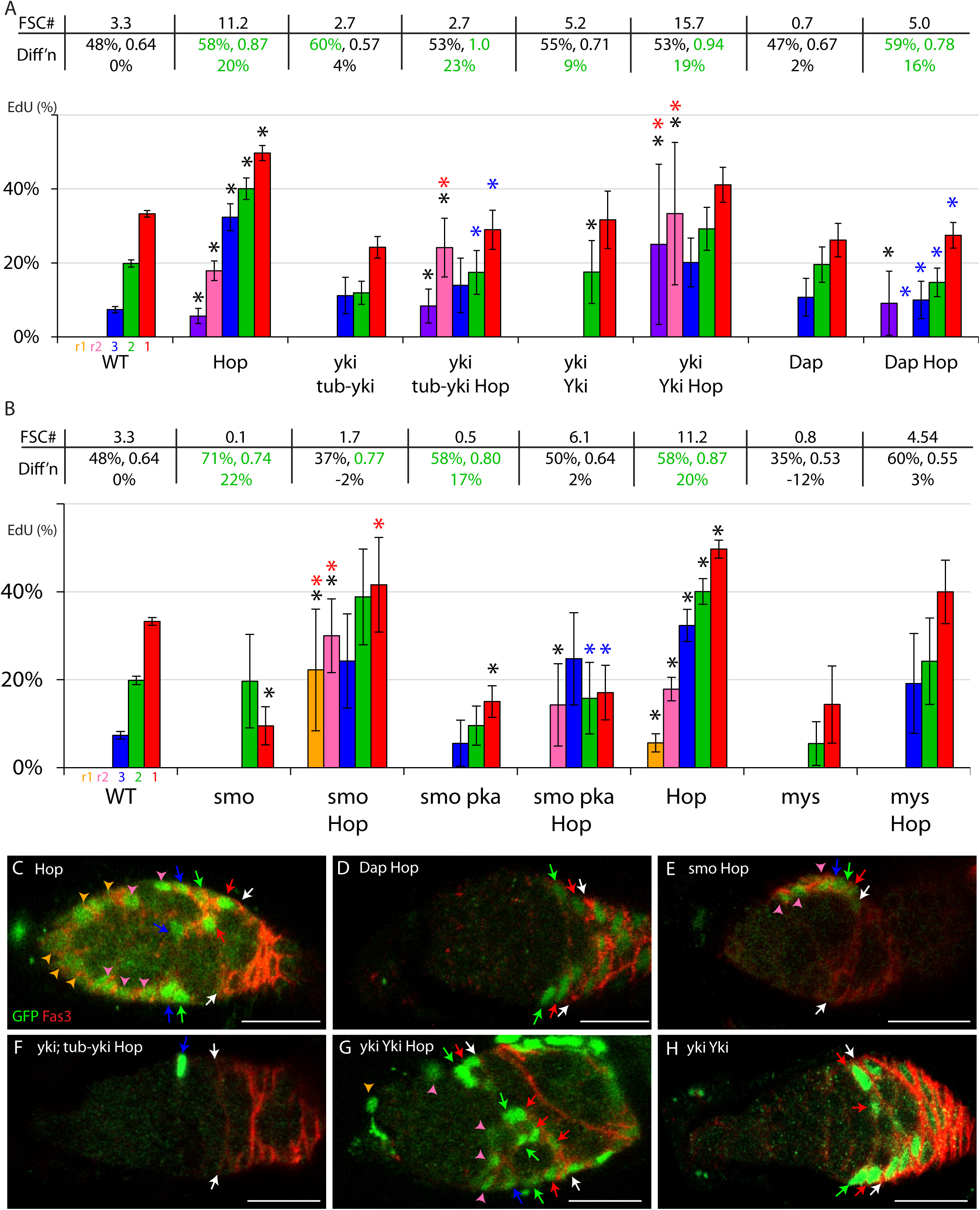
JAK-STAT effects are blocked by Cdk2 inhibition, partly depend on normal *yki* transcriptional regulation and largely epistatic to loss of Hh and integrin signaling. (A, B) Percentage of marked cells of the indicated genotypes (Hop, Yki, Dap indicate the named *UAS*-driven transgenes) that incorporated EdU for cells in the location of r1 ECs (purple), r2a ECs (pink), layer 3 (blue), layer 2 (green) and layer 1 (red) FSCs. SEMs and significant differences from control values are indicated (*, p<0.05, n-1 chi-squared test). Additional pair-wise comparisons show significant responses to a signal alteration in an altered candidate mediator background (red asterisk; *yki; tub-yki UAS-Hop* vs *yki; tub-yki*, *yki; UAS-Hop UAS-Yki* vs *yki; UAS-Yki*, *smo; UAS-Hop* vs *smo*, *smo pka: UAS-Hop* vs *smo pka*, *mys; UAS-Hop* vs *mys*) and whether the response is significantly lower than in a wild-type background (blue asterisks; *yki; tub-yki UAS-Hop* vs *UAS-Hop*, *yki; UAS-Hop UAS-Yki* vs *UAS-Hop*, *UAS-Dap UAS-Hop* vs *UAS-Hop*, *smo UAS-Hop* vs *UAS-Hop*, *smo pka UAS-Hop* vs *UAS-Hop*, *mys UAS-Hop* vs *UAS-Hop*). Absence of colored asterisks indicates no significant differences. For each genotype, above the graph, the average number of marked FSCs at 12d is indicated, as well as the percentage of FSCs in layer 1 at 6d, the calculated probability of FSC to FC conversion per cycle and (below those numbers) the inferred percentage loss of FSCs due to altered FC differentiation per 12h budding cycle. Large increases (green) or decreases (red) are highlighted. (C-H) MARCM samples for the indicated genotypes, with the Fas3 (red) border indicated (white arrows) and colored arrows indicating layer 1 (red), 2 (green) or 3 (blue) FSCs, r2a (pink arrowheads) and r1 (orange arrowheads) ECs. (C) The large increase of marked FSCs and derivatives in anterior locations due to excess JAK-STAT was (D) fully suppressed by excess Cdk2 inhibitor Dacapo. (E) Loss of FSCs due to loss of Hh signaling (*smo*) was partially suppressed by excess JAK-STAT. (F) Production of excess FSC derivatives by excess JAK-STAT was suppressed when *yki* was replaced by *tub-yki*. Substitution of *yki* with *UAS-Yki* also eliminates normal *yki* transcriptional control but likely leads to higher Yki levels; this (H) increased FSC numbers and (G) permitted further enhancement of FSC numbers by excess JAK-STAT. Scale bars 10μm.

EdU incorporation into cells in the EC region was stimulated by *UAS-Hop* almost as strongly in *yki; tub-yki* (15%) and *yki; UAS-yki* FSC derivatives (17%) as in a wild-type background (24%) (Fig. 9A). Since stimulation of division in the EC region requires both G1/S and G2/M stimulation, and the magnitudes of these responses exceed those due to *UAS-Stg* alone (which promotes G2 exit much more effectively than *UAS-Hop* according to FUCCI reporters), it appears that JAK-STAT can promote G1/S transitions effectively in EC territory without regulating *yki* transcription. These cells are described as being in EC territory rather than simply as ECs because they acquire several characteristics of FSCs and FCs when JAK-STAT pathway activity is elevated (18, 26).

Expression of the CycE/Cdk2 inhibitor, Dacapo (Dap) (34, 56), reduced FSC EdU incorporation only modestly in layer 1 (26%, 20%, 11% vs 33%, 20%, 7%) and did not significantly affect FC differentiation (Fig. 9A). FSC numbers were reduced more than expected (0.7 vs 3.3) from EdU index changes, suggesting that the EdU index may overestimate cycling frequency in this case (implying that S-phase may be lengthened). Complete loss of CycE fully arrests FSC division (21) and the *cycE^WX^* hypomorph, which retains an estimated 15% of normal CycE/Cdk2 kinase activity (21) reduced the EdU index (7%, 3%, 0) and FSC numbers (0.4) more than expression of Dap (data not shown), suggesting that Dap reduces CycE/Cdk2 activity by less than 80%. The EdU index of FSCs with excess Dap was not greatly altered by co-expression of *UAS-Hop* (to 27%, 15%, 10%). However, despite enhanced FC differentiation for layer 1 *UAS-Dap UAS-Hop* FSCs (0.78 vs 0.62), FSC numbers (5.0) were higher than for controls (Fig. 9A, D). The implication that division rate was significantly higher than indicated by EdU incorporation is consistent with direct deductions from live FUCCI reporters that S-phase was shortened by expression of *UAS-Hop*. Thus, reduction of CycE/Cdk2 kinase activity with Dap did not eliminate the proliferative response to increased JAK-STAT signaling but it reduced the response, whether measured by FSC numbers (5.0 vs 11.2) or FSC index (27%, 15%, 10% vs 50%, 40%, 32%). Stimulation of EdU incorporation in the EC region by *UAS-Hop* was also reduced by Dap (from 24% to 6%) (Fig. 9A). These results are consistent with the hypothesis that one of the actions of JAK-STAT signaling in FSCs and ECs is to raise CycE-Cdk2 activity for transitioning from G1 to S phase.

Both Hh and JAK-STAT pathways are major contributors to FSC cycling, raising the question of whether they might have complementary or overlapping mechanistic roles. Complete loss of Hh signaling (due to loss of *smo*) greatly reduced the EdU index and FSC numbers, while additional loss of PKA renders an intermediate level of constitutive pathway activity and a small increase in FSC survival (Fig. 9B). Increased JAK-STAT pathway activity significantly increased the EdU index of *smo* (from 10%, 20%, 0% to 42%, 39%, 24%) and *smo pka* mutant FSCs (from 15%, 10%, 6% to 17%, 16%, 25%), as well as the number of surviving *smo* (from 0.1 to 1.7) (Fig. 9E) and *smo pka* FSCs (from 0.5 to 6.1). The observed potential of excess JAK-STAT activity to compensate for reduced or absent Hh signaling is consistent with the prior deductions that Hh signaling acts primarily through *yki* induction, while JAK-STAT signaling acts partly through *yki* induction. Stimulation of EdU incorporation in EC region cells by *UAS-Hop* was not greatly affected in *smo* (28%) or *smo pka* FSC derivatives (14%). Previous studies suggest that the robust connection between Hh signaling and *yki* induction in FSCs may not be present in ECs (57), so ECs may not normally receive any proliferative input from Hh signaling, consistent with the unimpaired response of *smo* and *smo pka* genotypes to JAK-STAT.

Integrin signaling also normally stimulates FSC cycling through unknown mechanisms (58). Strong surface integrin staining is normally observed around FSCs but absent in more anterior regions where ECs reside (58). Increased JAK-STAT signaling induces ectopic integrin accumulation between FSC derivatives in the EC domain, suggesting that increased integrin signaling might underlie some part of the proliferative response to JAK-STAT signaling (18, 26). The FSC EdU index and FSC numbers were greatly reduced by partial inactivation of Myospheroid (Mys), the sole β-integrin expressed in Drosophila ovaries (59), (14%, 5%, 0%; 0.8 FSCs) but there was still a strong response to JAK-STAT in *mys; UAS-Hop* FSCs (EdU indices of 40%, 24%, 19%; 4.6 FSCs) (Fig. 9B). Thus, the proliferative actions of JAK-STAT in FSCs do not appear to depend significantly on increasing integrin signaling. The increased conversion of layer 1 FSCs to FCs, induced by increased JAK-STAT pathway activity alone and in combination with altered Yki, Dap (Fig. 9A) and Wnt pathway manipulations (18), was not evident in *mys* mutant FSCs, suggesting that integrin activity may play a role in this response (Fig. 9B).

There were only two marked *mys; UAS-Hop* FSC derivatives in EC territory, with no EdU incorporation, among samples with 77 marked FSCs. On average, expression of *UAS-Hop* alone produces about a third as many marked cells in the EC domain as FSCs (351/1041), much higher than controls (1085/5606= 19%) because cells in EC territory only proliferate in the former condition. The deficiency of marked *mys; UAS-Hop* cells in EC territory may therefore be an indication that division normally promoted by excess JAK-STAT is suppressed, but there also appears to be a reduction in anterior migration of FSCs, which prevents definitive evaluation of proliferative status. Thus, it is presently unclear, but possible that integrin induction is important for JAK-STAT to induce cell cycling in EC territory.

## Discussion

The regulation of adult stem cell proliferation is universally important to maintain stem cell populations while providing an adequate supply of derivatives to maintain tissues throughout life. For paradigms, like FSCs and mouse gut stem cells, where stem cell division and differentiation are independent, division and differentiation rates must be balanced over the whole population and mutations that accelerate division may often be critical to seeding cancers (4, 5). Additionally, for FSCs, there is pronounced spatial regulation of division rates, which serves to balance similarly graded differentiation rates and equalize stem cell potential for FSCs in different AP locations. Here, we have used FUCCI cell cycle reporters and epistasis analyses to gain new insights into the regulation of FSC division rate.

### Insights from FUCCI reporters

Live imaging with FUCCI reporters allowed us to record the frequency of cell-cycle transitions and deduce absolute cell cycle times. These parameters have not previously been measured for FSCs. In standard MARCM lineage experiments the total number of divisions of an FSC cannot be measured precisely because FC proliferation obscures the exact number of FCs derived directly from an FSC. Over a short period of time (like 3d), the number of FCs produced and FSC division rates can be estimated (4). However, this does not reveal the locations of FSC divisions and hence any information about the spatial pattern of FSC divisions. Here we found that layer 1 FSCs divide 3.4-fold faster than their anterior neighbors, roughly twice the difference previously estimated from EdU indices. The discrepancy arises from the deduction that S-phase is significantly longer in anterior than posterior FSCs (elevating anterior FSC EdU indices for a give rate of cycling).

For wild-type FSCs, the dramatic cycling differential between layer 1 and layer 2 FSCs, and the absolute cell cycle durations revealed by live FUCCI imaging revealed that the replenishment of posterior (layer 1) FSCs suffices to match conversion to FCs, and hence that there is little or no net flow from anterior to posterior within the FSC domain. Live imaging measurements also resolved a minor conundrum with regard to spatial regulators of FSC division. It was previously found that JAK-STAT signaling is a major contributor to FSC division in all locations. The FSC EdU index remained spatially graded in the absence of STAT activity, clearly indicating another source of AP pattern (18). By contrast, supplying excess JAK-STAT activity in a pattern complementary to the normal posterior to anterior gradient, thereby equalizing activity over the FSC domain, resulted in almost identical EdU indices among the three FSC layers (18). Here we found that under those conditions, layer 1 and layer 2 FSC FUCCI profiles remained very different. Moreover, the overall cell cycle duration ratio (layer 2: layer 1) declined, but only from 3.4 to 2.6; EdU indices were co-incidentally equal because S-phase remained substantially longer in anterior than posterior FSCs. Thus, FUCCI reporter tests now clearly show a major spatial influence on FSC cycling under conditions of both STAT absence and uniform JAK-STAT activity. This unknown second influence supplements the effects of graded JAK-STAT signaling on cell division rate under normal conditions.

Results of H2B-RFP dilution experiments also supported the general conclusion of faster cycling of layer 1 than layer 2 FSCs, with very little cell division further anterior. This approach had some systematic limitations that precluded accurate measurement of division rates of cells in a specific location. First, it did not prove possible to achieve highly similar H2B-RFP labeling intensities among cells in a given AP location prior to chase. Second, it is known from earlier studies that FSCs can change AP location quite frequently. So, for example, an FSC scored in layer 2 after several days of chase may have resided in layer 1 or layer 3 for some of that time. Consistent with this behavior, there were sometimes dramatic differences of H2B-RFP signal among FSCs in a given layer. Nevertheless, it was clear that H2B-RFP dilution was fastest in FCs and faster in layer 1 than in more anterior FSCs. We also used live imaging of samples after significant periods of H2B-RFP dilution to visualize cell movements directly. H2B-RFP intensities allowed us to define FSC layers and track individual cells with dilutions (and hence, division histories), characteristic of their location. This showed for the first time that FSCs can move in either direction between layers, consistent with the deduction that there is little net flow within the FSC domain.

We also gained some information from FUCCI reporters regarding the specific cell cycle transitions spatially regulated by signals. Reduced JAK-STAT activity in layer 1 FSCs reduced S-phase representation and increased G2-phase frequency, suggesting that JAK-STAT normally promotes both G1/S and G2/M transitions. This deduction is consistent with the effects of increased JAK-STAT activity in each FSC layer. Here, G2 frequency was reduced, clearly indicating G2-M stimulation. The G1:S ratio actually increased, but we still infer facilitation of a G1/S transition by comparing to the effects of excess Stg (expected only to directly stimulate G2-M passage), which results in much higher G1, and lower S-phase frequencies. Similar logic suggested that the PI3K pathway also may promote both transitions but with a lesser normal role than JAK-STAT for the G1/S transition. The promotion of a G1/S transition is also evident for excess JAK-STAT in the most anterior ECs, which normally reside in G1, whereas excess PI3K was without effect.

The deduction of primary actions of regulators through FUCCI reporter responses has significant limitations, especially if examined only in fixed samples and when testing potentially non-physiological excess activities. Excess CycE is expected to accelerate the G1/S transition. The G1 proportion in fixed samples and G1 duration in live imaging were indeed both reduced. However, live imaging revealed that G2 was also shortened and S-phase was lengthened, neither of which could be deduced just from fixed samples. Moreover, it is unlikely that those are direct effects of excess CycE. In wild-type FSCs there is spatial regulation of the duration of S-phase in addition to G1 and G2, the more commonly studied targets of regulation. Since phases are cyclical, with precise internal sequences of events, and prior phases prepare molecular conditions for future events (35, 60–62), it is possible that the variation in S-phase relates to differences in preparation rather than signaling inputs that act during S-phase.

With these caveats in mind, we can still deduce some of the likely actions of the presently unknown spatial regulator(s) of FSC division. In germaria with near-uniform JAK-STAT signaling, the biggest fractional difference in cell cycle phase duration between layer 2 and layer 1 FSCs is for G1 (548 vs 148 mins; G2 is 661 vs 303min; S is 623 vs 237 min). Passage out of G1 is also clearly the biggest limitation for ECs under these conditions. The simplest hypothesis is that there is a regulatory signal that is very strong in ECs, and much stronger in anterior than posterior FSCs, which primarily restricts G1/S passage. Although we favor the concept of complementary distributions of stimulatory (JAK-STAT) and inhibitory signals, the missing signal could alternatively be stimulatory and posteriorly-biased.

### Exploration of signals and signal mediators

The Hpo/Yki pathway is a commonly employed regulator of cell growth, which can intersect with several signaling pathways and respond to a variety of mechanical cues (29). Earlier studies revealed a major positive role of Yki activity in FSC proliferation and competitive survival, including evidence that *yki* transcription was regulated by Hh signaling and responsible for the proliferative effects of Hh (17). Those deductions were broadly supported here, now that we are able to monitor all FSCs (10). Indeed, FSC division, monitored by EdU index in MARCM tests, was drastically reduced in *yki* and *smo* mutant FSCs, but fully restored to *smo* mutant FSCs by excess activated Yki. Moreover, as found previously (17), the increase in FSC numbers and EdU index induced by *ptc* mutation (activating the Hh pathway fully) was completely suppressed by substituting *yki* with *tub-yki* to eliminate normal transcriptional control of *yki*. However, the magnitudes of EdU increases we observed for *ptc* and loss of inhibitors of Yki activity (*hpo*, *kibra*, *wts*) were much lower than reported previously and were mostly restricted to more anterior FSCs (the subset of FSCs scored previously was likely an anterior subset). Here, we were also able to measure other parameters of FSC behavior. We found no significant changes in FSC differentiation to FCs for *ptc*, *hpo*, *kibra* and *wts* genotypes and similar moderate increases in the number of marked FSCs over time. These data support a conclusion that these genetic alterations primarily induce a small increase in cycling in anterior FSCs, sufficient to enhance competitive survival. Plausibly, this is because Yki and Hh pathway activity are already high in normal FSCs, as supported by prior observation of the *ptc-lacZ* Hh pathway reporter (25, 26). That level of activity was roughly reproduced in FSCs of the genotype *smo pka; Su(fu)* (*26*). Those FSCs cannot respond to Hh and were found to produce a normal AP EdU index profile. All of these factors support the conclusion that the shallow anterior to posterior gradient of Hh signaling across the FSC domain does not have a major impact on the spatial pattern of FSC divisions.

Removing transcriptional control of *yki* also reduced the stimulation of EdU incorporation by increased JAK-STAT activity when replacing *yki* function with either *tub-yki* or *UAS-Yki*. In the latter case, there was still a significant response to increased JAK-STAT, suggesting that JAK-STAT FSC responses are only partly dependent on induction of *yki*. Stimulation of EdU incorporation into ECs by excess JAK-STAT was unaffected by these manipulations, providing further evidence of some stimulatory actions of JAK-STAT unrelated to transcriptional regulation of *yki*. Thus, while JAK-STAT must have additional mediators, transcriptional regulation of *yki* may be a point of convergence for proliferative FSC signals emanating from the anterior (Hh) and posterior (JAK-STAT ligand). Epistasis results with excess PI3K also suggested that FSC responses were partly dependent on *yki* transcriptional regulation. However, it will be important to explore PI3K responses more fully because increased EdU indices induced by PI3K were generally accompanied by reduced FSC numbers, for which we currently have no sound explanation.

### Comparison to other stem cell paradigms

Mouse gut and epidermal stem cells share with FSCs the characteristics of generally high constitutive division rates and division-independent differentiation (5, 6). Despite the acknowledgment that division and differentiation can be regulated independently, the majority of studies assessing contributions of specific signals measure stem cell survival or amplification, rather than the individual contributing parameters. Nevertheless, there is evidence of a gradient of Wnt pathway activity in gut crypts (63) and increased division rates were evident in organoids when pathway activity was increased by *Apc* mutation (64). Division stimulation through the EGFR family was inferred from increased phospho-histone incorporation when the Lrig1 negative feedback regulator was removed (65), while an effect of Notch signaling was inferred from effects on expression of Cdk inhibitors (66). A positive role of Yki orthologs YAP and TAZ, and antagonism by the Hippo pathway was deduced from EdU incorporation and Ki67 proliferation marker staining (67). Similar methods were used to deduce that loss of the mitochondrial pyruvate carrier stimulates stem cell division (68). None of these studies, however, includes a quantitative measure of stem cell cycling rates *in vivo*, and deductions rely on uncertain assumptions of constant S-phase length or Ki67 marker interpretation (69). Since mouse gut crypts can be imaged over long periods of time *in situ* (5), these limitations might be resolved by using live FUCCI reporter imaging in the same way we employed for FSCs.

In the basal epidermal stem cell layer there is indirect evidence of ECM-integrin interactions supporting proliferation (70). Live imaging showed that division is generally consequent to departure of a neighboring differentiating cell from the basal layer (71, 72), while other studies implicate membrane tension of differentiated neighbors, potentially transmitted mechanically (73). Thus, stem cell division rate may largely be regulated indirectly in this paradigm, consequent to non-cell-autonomous cues from differentiating neighbors.

In the male Drosophila gonad, the cell type closest to female FSCs are somatic Cyst stem cells (CysSCs). They share the properties of maintenance by population asymmetry and being more proliferative than immediate neighbors, hub cells (74). However, their differentiated daughters, in contrast to FCs, do not divide and their overall function guiding germline differentiation combines EC and FC functions. Both Hh and JAK-STAT signaling promote CysSC division, apparently through indirect induction of Hippo pathway components (75). Other studies suggest that loss of upstream Hippo pathway regulators, Merlin and Expanded may promote CysSC division via increased MAPK and PI3K pathway activities (76). CysSC cycling was also found to be stimulated by activin receptor signaling, with the presence of Follistatin preventing an analogous response in hub cells (77).

It is clear that there are huge gaps in quantitative measurements of stem cell cycling, relevant signals and their mediators for even the best-studied examples of population asymmetry, where stem cell competition and communal function depend critically on regulation of division rates. Our considerable knowledge of relevant FSC signals, the potential to test causal relationships genetically and the potential to measure cell cycling quantitatively through FUCCI reporters and live imaging, demonstrated here, make FSCs an especially promising paradigm for understanding the regulation of stem cell division rates. Other stem cells, including neural and muscle stem cells, have been studied in the notably different context of transitioning into or out of quiescent states (23, 78, 79). Such studies are beginning to reveal a diversity of mechanisms, including clear evidence that the G2/M transition can be regulated, supplementing the long-held assumption that regulation was primarily through entering a G0 state prior to S-phase (24, 80).

## Acknowledgments

This work was supported by NIH RO1 GM079351 to DK. We thank Burcu Gulez for contributions to H2B-RFP studies, Bruce Edgar (University of Washington, Seattle), Laura Buttitta (University of Michigan, Ann Arbor), Bob Duronio (University of North Carolina, Chapel Hill), Carl Thummel (University of Utah, Salt Lake City), Alana O’Reilly (Fox Chase Cancer Center, Philadelphia) and Laura Johnston (Columbia University, New York) for reagents, the Bloomington stock center for provision of genetic reagents, the Developmental Studies Hybridoma Bank (DSHB) for antibodies, FlyBase as an information resource, and the confocal microscope resource provided by the Dept. of Biological Sciences, Columbia University.

## Experimental Procedures

### FUCCI Reporter Experiments

Nuclear-targeted GFP and RFP UAS-driven FUCCI reporters on the second (BL-55121) and third (BL-55122) chromosomes were combined and expressed conditionally using *C587-GAL4* and a second chromosome temperature-sensitive *GAL80* transgene (from BL-7108). 1-3d old female flies of the genotype *C587-GAL4; UAS-FUCCI/tub-ts-GAL80, FRT42D tub-lacZ; (UAS-X)/UAS-FUCCI* were collected, where *UAS-X* was absent (wild-type), *UAS-CycE*, *UAS-Stg* (BL-4778), *UAS-Stg* + *UAS-CycE*, *UAS-Hop^3W^, UAS-Hop^3W^ + UAS-Dap, UAS-dnTCF* (BL-4785), *UAS-dnTCF* + *UAS-Hop*, *UAS-stat RNAi* (BL-31317) + *UAS-DIAP1*, *UAS-PI3K92E A2860C* (aka Dp110-DN) (BL-8289) or *UAS-PI3K92E* (BL-8287) (transgenes not specified were the same as in (18)). Flies were incubated at 29C for 3d to inactivate GAL80. Dissected ovaries underwent the EdU and immunohistochemistry protocols described below, without staining for GFP.

### Live Imaging

Imaging chambers were fabricated as described previously (10, 46). Ovaries from flies with *C587-GAL4*, *UAS-FUCCI* genotypes described above and incubated at 29C for 3d were dissected into imaging medium formulated as in (81) (20% FBS in Schneiders insect medium, 0.2 mg/mL insulin, penicillin and streptomycin). After separating ovaries into individual ovarioles, 135 µL ovarioles were mixed with 15 µL Matrigel (Corning), added to the imaging chamber and left covered for 15 min to gel. Wells were then filled to the top with imaging medium. Germaria were generally imaged every 15 min on a Zeiss LSM 700 confocal microscope.

### MARCM Clonal Analysis

1-3d old adult *Drosophila melanogaster* females with the appropriate genotypes were given a single 30 min (for *FRT40A* and *FRT42D*) or 45 min (for *FRT82B*) heat shock at 37C. Afterwards, flies were incubated at 25C, with the exception of experiments using *FRT40A* or *FRT42D* and *UAS-Hop*, where 29C was used to increase GAL4 activity sufficiently to induce a strong increase in JAK-STAT pathway activity, exactly as in prior analyses (18). Flies were maintained by frequent passage on normal rich food supplemented by fresh wet yeast during the 12d experimental period. Flies were dissected at 6d and 12d. Immediately after dissection, 6d ovaries underwent 1h of EdU labelling based on the protocol of the Click-iT™ Plus EdU Cell Proliferation Kit for Imaging (Invitrogen). Both 6d and 12d ovaries were stained for Fasciclin III (Fas3) and GFP. Ovaries were then manually separated into constituent ovarioles, and mounted using DAPI Fluoromount-G® (SouthernBiotech) to stain nuclei. Ovarioles were imaged with a Zeiss LSM700 or LSM800 confocal microscope, operated in part by the Zeiss ZEN software.

The entire germarium was captured in the images, as well as an average of 3-4 egg chambers. Collected images were saved as CZI files, and were later analyzed utilizing the ZEN Lite software. We aimed to image at least 50 germaria for every genotype in each experiment.

### MARCM Genotypes

Flies with alleles on an *FRT19A*, *FRT40A*, *FRT42D*, or *FRT82B* chromosomes were used in MARCM experiments using the following genotypes:

FRT40A: *yw hs-Flp, UAS-nGFP, tub-GAL4 /yw; act-GAL80 FRT40A / (X)FRT40A; act>CD2>GAL4/ Y* – where X, Y combinations included: (X) – *NM* (Nuclear Myc, Control), *cycE^AR95^*, *smo^2^*, *smo^2^ pka^B3^*, *chico^1^*, *pten^c494^* (Y) – *UAS-CycE*, *UAS-Stg*, *UAS-CycE* + *UAS-Stg*, *UAS-Yki* (BL-28819), *UAS-YkiS168A* (BL-28818), *UAS-Hop^3W^, UAS-Dap* (BL-83334)*, UAS-PI3K*, *UAS-PI3K* + *UAS-CycE*, *UAS-CycD* + *UAS-Cdk4* (Dr. Bruce Edgar), or *Su(fu)^LP^* (also present on the *act>CD2>GAL4* chromosome in this experiment).

FRT42D: *yw hs-Flp, UAS-nGFP, tub-GAL4 /yw; FRT42D act-GAL80 tub-GAL80 / FRT42D (X); act>CD2>GAL4/ Y* – where X, Y combinations included: (X) – *sha* (Control), *ubi-GFP* (Control), *arr^2^, yki^B5^*, *hpo^42-47^*, *ptc^S2^*, *ptc^S2^ yki^B3^*, (Y) – *tub-yki* (17), *tub-yki* +*UAS-Hop^3W^, UAS-Yki* +*UAS-Hop^3W^, UAS-PI3K* + *UAS-Yki*, *tub-yki* +*UAS-Hop^3W^*, or *UAS-Dap* +*UAS-Hop^3W^*.

FRT82B: *yw hs-Flp, UAS-nGFP, tub-GAL4 /yw; act>CD2>GAL4 UAS-GFP / Y; FRT82B tub-GAL80/FRT82B (X)* – where X, Y combinations included: (X) – *NM* (control), *stat^85C9^, kibra^32^, wts^x1^, kibra^32^* + *stat^85C9^*, *UAS-CycE* + *stat^85C9^*, *UAS-CycE* + *kibra^32^* + *stat^85C9^*, *dMCP1^1^*(BL-83685), (Y) *UAS-Stg*.

FRT19A: hs-flp tub-GAL80 FRT19A / (X) FRT19A; act-GAL4, UAS-GFP / Y-where X was *+* (wild-type) or *mys^n12^*, and Y was *+* (wild-type) or *UAS-Hop^3W^*.

### EdU Protocol

Ovaries were directly dissected into a solution of 15 µM EdU in Schneider’s *Drosophila* media (500µl, Gibco) and incubated for one hour at room temperature. These tubes were laid on their side and rocked manually, to ensure all dissected ovaries were fully submerged. Ovaries were then fixed in 3.7% paraformaldehyde in PBS for 10 minutes, treated with Triton in PBS (500 µl, 0.5% v/v) for 20 minutes, and rinsed 2x with bovine serum albumin (BSA) in PBS (500 µl, 3% w/v) for 5 minutes each rinse. Ovaries were exposed to the Click-iT Plus reaction cocktail (500 µl) for EdU visualization, for 45 minutes. The reaction cocktail was freshly prepared prior to use, with reagents from the Invitrogen™ Click-iT™ Plus EdU Cell Proliferation Kit for Imaging, including the Alexa Fluor™ 594 dye. Ovaries were then rinsed 3x with BSA in PBS (500 µl, 3% w/v) for 5 minutes each rinse.

### Immunohistochemistry

For experiments without EdU, ovaries were dissected directly into a fixation solution of 4% paraformaldehyde in PBS for 10 min at room temperature, rinsed 3x in PBS, and blocked in 10% normal goat serum (NGS) (Jackson ImmunoResearch Laboratories) in PBS with 0.1% Triton and 0.05% Tween-20 (PBST) for 1 h. Monoclonal antibodies for Fas-3 were obtained from the Developmental Studies Hybridoma Bank, created by the NICHD of the NIH and maintained at The University of Iowa, Department of Biology, Iowa City, IA 52242. 7G10 anti-Fasciclin III was deposited to the DSHB by Goodman, C. and was used at 1:250 in PBST. Other primary antibodies used were anti-GFP (A6455, Molecular Probes) at 1:1000 in PBST. Ovaries were incubated in primary antibodies overnight, rinsed three times in PBST, and incubated 1-2 h in secondary antibodies Alexa-488 and Alexa-647 (ThermoFisher) at 1:1000 in PBST to label GFP and Fas3, respectively. DAPI-Fluoromount-G (Southern Biotech) was used to mount ovaries.

### Imaging and scoring

All germaria were imaged in three dimensions on an LSM700 or LSM800 confocal laser scanning microscope (Zeiss) and using a 63x 1.4 N.A. lens. Zeiss ZEN software was used to operate the microscope and view images. Images were typically 700×700 pixels with a bit depth of 12. The scaling per pixel was 0.21 µm x 0.21 µm x 2.5 µm. The range indicator in ZEN was used to determine the appropriate laser intensity and gain. ZEN was used to linearly adjust channel intensity for dim signals to improve brightness without photobleaching samples. Images were saved as CZI files and scored directly in ZEN. DAPI and Fas3 staining were used as landmarks to guide scoring. Marked cells were considered FSCs if they were within three cell diameters anterior of the Fas3 border. Cells immediately adjacent to the border were considered to be in Layer 1, with Layers 2 and 3 in sequentially anterior positions. Anterior to the FSC niche, the EC region was roughly divided into two halves, with region 2a ECs immediately anterior to FSCs and region 1 ECs anterior to that. Germaria were also scored (Y/N) for the presence of marked FCs. For the “Immediate FC Method” (18), the presence of an FC immediately posterior to Layer 1 was also scored Y/N. For publication, images were digitally zoomed in ZEN and exported as *tif* files using the “Contents of Image Window” function. Images were rotated in Abode Photoshop CS5 to uniformly orient the germaria.

### H2B-RFP Dilution Experiments

The original third chromosome P-element insertion of mRFP N-terminally tagged H2B as *UAS-H2B-RFP* in a mini[*w^+^*] vector is described in (82). Transposon mobilization, as in (83), was used to identify new insertion sites based on altered eye color and H2B-RFP expression of new lines was assessed after crossing to an *act>GAL4* driver. Parent flies of *genotypes yw hs-flp; tub-GAL80(ts) FRT42D tub-lacZ / CyO; act>GAL4, UAS-GFP / TM2* and *yw; FRT42D / CyO; UAS-H2B-RFP/ TM2* were crossed at 18C to produce female progeny of the experimental genotype *yw hs-flp / yw; tub-GAL80(ts) FRT42D tub-lacZ / FRT42D; act>GAL4, UAS-GFP / UAS-H2B-RFP*. Crosses or adult progeny were shifted to 29C for various times to allow H2B-RFP expression and then back to 18C to prevent further H2B-RFP expression. Flies were regularly changed to new vials with added fresh moist yeast. Ovaries were dissected and stained for Fas3 and Traffic Jam (using guinea pig antibody from Dr. Dorothea Godt). Traffic Jam staining was used to outline nuclei using the Draw Spline Contour function for measuring H2B-RFP intensity with Zen software. For live imaging the genotype of flies was *yw hs-flp; ubi-NLS-GFP FRT40A / tub-GAL80(ts) FRT42D tub-lacZ; act>GAL4, UAS-GFP / UAS-H2B-RFP* so that all nuclei were marked by GFP. Live imaging was performed as described above but with 5 min intervals to track cell movements.

### Statistics and Reproducibility

All images shown are representative of at least ten examples. No statistical method was used to predetermine sample size but we used prior experience to establish minimal sample sizes. No samples were excluded from analysis, provided staining was of high quality (exclusion of live imaging FUCCI samples with no cell-cycle phase transitions is described in Results). Investigators were not blinded during outcome assessment, but had no pre-conception of what the outcomes might be. For MARCM studies, the EdU index was calculated for marked FSCs in each layer, for r1 and r2a ECs. The “N-1” Chi-squared test method was used to calculate a Z score for determining significance of any differences between indicated genotypes for cells in the same location, and error was reported as standard error of a proportion. To determine whether the EdU index distribution among the FSC layers of an altered genotype differed significantly from controls, we first calculated the average EdU index for all FSCs of the altered genotype, with each layer contribution weighted based on the normal distribution of FSC among layers measured in appropriate controls. This average EdU index was then multiplied by the control EdU index for each layer to derive expected EdU indexes for each layer of an altered genotype if the EdU pattern matched controls. Finally, a chi-squared test was applied to compare observed and expected EdU indexes for each layer to determine the statistical significance of differences. Graphs of EdU indices also include tabulation of the average number of marked FSCs at 12d, the percentage of marked FSCs present in layer 1, and the probability (p) of a marked layer 1 FSC becoming an FC in a single 12h egg chamber budding cycle, calculated as previously documented (18). From the latter two parameters, we calculated the percentage change in conversion of FSCs to FCs relative to wild-type per budding cycle, as described below.

The loss of a marked FSC of a specific genotype by differentiation to an FC each cycle, F = f (fraction of FSCs in layer 1) x p(differentiation to FC) per marked FSC.

For WT FSCs, F = 0.48 x 0.64 = 0.307

The change in FSC conversion to FCs per FSC per cycle for a different FSC genotype, ΔF = (fp −0.307).

### Data Availability

All data supporting the findings from this study are available from the corresponding author upon reasonable request.

