## Supplementary Figure 1 for "Spatial regulation of Drosophila ovarian Follicle Stem Cell division rates and cell cycle transitions"

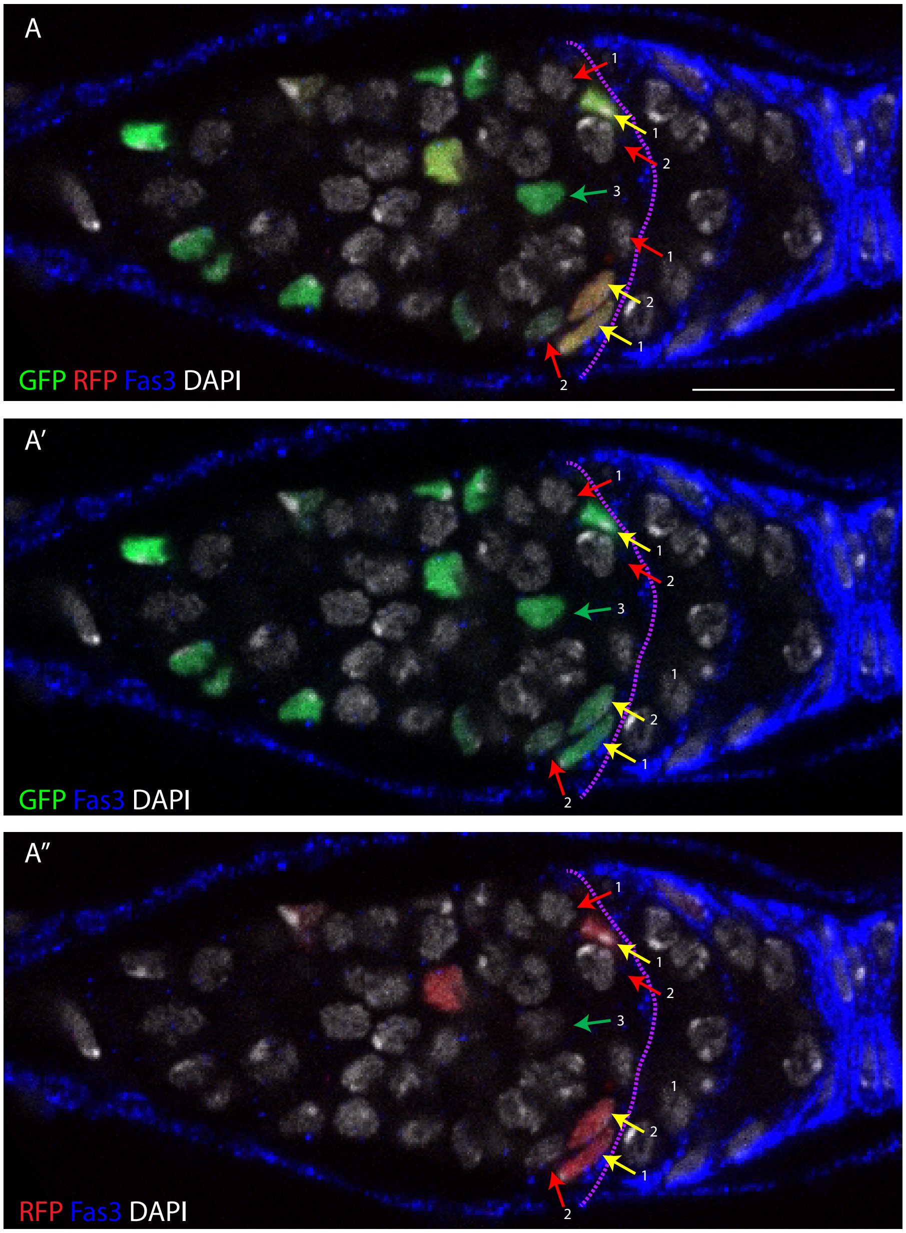


**Figure S1. Scoring cell cycle phases using FUCCI reporters and DAPI staining without EdU.**

A single z-section of a *C587>FUCCI* germarium stained for DAPI (white) to show all nuclei. The anterior border of strong Fas3 (blue) expression is outlined by a purple dotted line, allowing designation of FSCs in different layers (labeled as 1, 2 or 3). FSCs in G1 (green arrows; GFP only), G2 (yellow arrows; GFP and RFP) or S phase (red arrows; no GFP or RFP) are indicated. Scale bar 10μm.
